# A Statistical Procedure for Genome-wide Detection of QTL Hotspots Using Public Databases with Application to Rice

**DOI:** 10.1101/479725

**Authors:** Man-Hsia Yang, Dong-Hong Wu, Chen-Hung Kao

## Abstract

Genome-wide detection of quantitative trait loci (QTL) hotspots underlying variation in many molecular and phenotypic traits has been a key step in various biological studies since the QTL hotspots are highly informative and can be linked to the genes for the quantitative traits. Several statistical methods have been proposed to detect QTL hotspots. These hotspot detection methods rely heavily on permutation tests performed on summarized QTL data or individual-level data (with genotypes and phenotypes) from the genetical genomics experiments. In this article, we propose a statistical procedure for QTL hotspot detection by using the summarized QTL (interval) data collected in public web-accessible databases. First, a simple statistical method based on the uniform distribution is derived to convert the QTL interval data into the expected QTL frequency (EQF) matrix. And then, to account for the correlation structure among traits, the QTLs for correlated traits are grouped together into the same categories to form a reduced EQF matrix. Furthermore, a permutation algorithm on the EQF elements or on the QTL intervals is developed to compute a sliding scale of EQF thresholds, ranging from strict to liberal, for assessing the significance of QTL hotspots. With grouping, much stricter thresholds can be obtained to avoid the detection of spurious hotspots. Real example analysis and simulation study are carried out to illustrate our procedure, evaluate the performances and compare with other methods. It shows that our procedure can control the genome-wide error rates at the target levels, provide appropriate thresholds for correlated data and is comparable to the methods using individual-level data in hotspot detection. Depending on the thresholds used, more than 100 hotspots are detected in GRAMENE rice database. We also perform a genome-wide comparative analysis of the detected hotspots and the known genes collected in the Rice Q-TARO database. The comparative analysis reveals that the hotspots and genes are conformable in the sense that they co-localize closely and are functionally related to relevant traits. Our statistical procedure can provide a framework for exploring the networks among QTL hotspots, genes and quantitative traits in biological studies. The R codes that produce both numerical and graphical outputs of QTL hotspot detection in the genome are available on the worldwide web http://www.stat.sinica.edu.tw/~chkao/.

## INTRODUCTION

Quantitative trait loci (QTL) detection has been a key step to provide deeper insight into the genetic mechanism of quantitative traits in many areas of biological researches, including crops, evolution, ecology and genetical genomics studies etc. (Lander and Botstein 1989; Haley and Knott 1992; Jansen 1993; Zeng 1994; Kao *et al.* 1999; Sen and Churchill 2001; Broman 2003; Kao 2006; Lee *et al.* 2014; Wei and Xu 2016). In QTL detection, it is often found that some of the genomic regions are relatively enriched in QTLs as compared to other regions, and that QTLs responsible for correlated traits frequently co-localize in some specific genetic regions (Goffinet and Gerber 2000; Schadt *et al.* 2003; Chardon *et al.* 2004; West *et al.* 2007; Breitling *et al.* 2008; Wu *et al.* 2008; Li *et al.* 2010; Ali *et al.* 2013; Basnet *et al.* 2015). The regions enriched in QTLs are usually called QTL hotspots, and, statistically, they harbor a significantly higher number of QTLs than expected by random chance. There are several possible reasons for the phenomenon of QTL hotspots: Firstly, QTLs explaining most trait variations can be effectively and consistently detected and mapped to similar regions across different experiments in various studies. Secondly, QTLs with higher allelic diversity have a greater chance of being detected in different crosses and environments (Zhao *et al.* 2011; Vuong *et al.* 2015; Mengistu *et al.* 2016). Thirdly, pleiotropic or closely linked QTLs for correlated traits (Falconer and Mackay 1996) will be frequently detected at the same regions in different experiments. For example, a gene called *SCM2/ APO1* in rice exhibits effects on panicle structure, culm strength and lodging resistance (Ookawa *et al.* 2010), and a gene called *GST* in maize shows the resistance to northern leaf blight, southern leaf blight and gray leaf spot diseases (Wisser *et al.* 2011; Ali *et al.* 2013). A genetical genomics study of *Arabidopsis*, Fu *et al.* (2009) found that the detected hotspots can be linked to several well-studied genes with pleiotropic effects on plant metabolism, physiology and morphology and development. Therefore, hotspot detection can lead to identifying genes that affect the relevant traits (Chardon *et al.* 2004; Fu *et al.* 2009). As the QTL hotspots are highly informative and may harbor genes for the target traits, the detection of QTL hotspots at the genome-wide level has been an important task in broad areas of biological studies (Breitling *et al.* 2008; Fu *et al.* 2009; Neto *et al.* 2012; Frary *et al.* 2014).

Genome-wide detection of QTL hotspots requires the collection of many QTLs for numerous and widespread traits d in the genome to enable the detection analysis. Genetical genomics experiments and public QTL databases are two feasible ways to provide such data with numerous QTLs for genome-wide QTL hotspot detection. A single genetical genomics experiment can produce abundant individual-level data containing the original genotypes (genetic markers) and thousands of molecular traits (phenotypic traits), such as gene expressions or protein contents, in a single segregation population. By treating the molecular traits as quantitative traits, the QTL mapping procedure can be performed to detect QTLs by providing the LOD scores at every genomic position for each trait. The LOD scores at every position for all traits can be recorded into a LOD score matrix, and given a predetermined LOD threshold, the LOD score matrix can then be converted into a QTL matrix by assigning 1 to the detected QTL positions and 0 otherwise. Using a genetical genomics experiment, West *et al.* (2007), Wu *et al.* (2008) and Li *et al.* (2010) permuted the QTL matrix across the genomic positions separately by traits to generate null distribution of hotspot sizes and compute the thresholds for assessing the significance of QTL hotspots. As these methods do not account for the correlation structure among traits, the hotspot size thresholds are severely underestimated, leading to the detection of too many spurious hotspots (Breitling *et al.* 2008). To consider the correlation structure among traits, Breitling *et al.* (2008) permuted the individual-level data by swapping the phenotypes between individuals and keeping the genotypes intact to generate the permuted data sets, and then performed QTL mapping on all the permuted data sets to obtain the QTL matrices for determining the thresholds. The method by Breitling *et al.* (2008) can overcome the underestimation of thresholds in the hotspot detection, but may neglect small and moderate hotspots with strong LOD scores as the magnitude of LOD score is not considered (Neto *et al.* 2012). To consider the magnitude of LOD score, Neto *et al.* (2012) adopted the same permutation and QTL mapping schemes as in Breitling *et al.* (2008) to obtain LOD score matrices. The LOD score matrices are then used to determine a sliding scale of empirical LOD thresholds given a range of possible spurious hotspot sizes in assessing the significance of QTL hotspots. In this way, the approach of Neto *et al.* (2012) can effectively discover small and moderate hotspots with strong LOD scores.

Besides using genetical genomics experiments, using public databases is also an effective and convenient way to obtain genome-wide detection of QTL hotspots. Several public databases, such as GRAMENE (http://www.GRAMENE.org/), Q-TARO (http://qtaro.abr.affrc.go.jp/), Rice TOGO browser (http://agri-trait.dna.affrc.go.jp/index.html), PeanutBase (http://peanutbase.org) and MaizeGDB, contain diversity in traits and wide-ranging distribution in the genome from numerous independent QTL mapping experiments, thus they can serve as an alternative source of genome-wide QTL hotspot detection. In these public databases, only the flanking markers of the detected QTLs (the QTL intervals), trait names and reference sources are curated, and no individual-level data is available to allow the hotspot analyses of Breitling *et al.* (2008) and Neto *et al.* (2012). Chardon *et al.* (2004) self-collected QTL data from the literature and developed a statistical method based on the normal distribution for hotspot detection. As the method of Chardon *et al.* (2004) requires the estimates of QTL positions and their variances in computation, it is not applicable to the above mentioned public databases, either. In this article, by using public databases, we combine and modify the key ideas of the above-mentioned methods to propose a statistical procedure for QTL hotspot detection at the genome-wide level. We first develop a statistical method based on the uniform distribution to convert the QTL intervals into the expected QTL frequency (EQF) matrix. Then, to cope with the correlation structure among the traits, we group the correlated traits into the same trait categories to form a reduced EQF matrix. Furthermore, inspired by the works of Neto *et al.* (2012) and Cabrera *et al.* (2012), we devise a permutation algorithm on the EQF elements (bins) as well as on the QTL intervals to compute a sliding scale of hotspot size thresholds given a range of possible spurious hotspot numbers for assessing the QTL hotspots. In this way, our statistical procedure can effectively correct the underestimation of threshold and result in less spurious hotspots. Simulation study shows that our statistical procedure can control the genome-wide error rates at the target levels, can provide appropriate thresholds for correlated data, and is comparable to the methods using individual-level data in hotspot detection. In real example analysis, the 8216 QTLs responsible for 236 different traits from 230 independent worldwide studies in GRAMENE rice database are analyzed for hotspot detection. The detected hotspots are compared with the 122 known genes collected in Q-TARO rice database to explore the interplays among QTL hotspots, genes and quantitative traits. The R codes that produce both numerical and graphical summaries of the QTL hotspot detection in the genomes are available on the worldwide web http://www.stat.sinica.edu.tw/~chkao/. Our analyses can establish a framework for exploring the networks among QTLs, genes and traits in biological studies.

## STATISTICAL METHODS

Our proposed procedure operates on the QTL intervals and can take the correlation structure among traits into account for QTL hotspot detection. In the following, the QTL intervals in the GRAMENE database that can be integrated into a QTL matrix is first described. Then we propose a simple method based on the uniform distribution to convert the QTL matrix into the EQF matrix and further into the reduced EQF matrix by grouping of correlated traits into the same trait categories. After that, we develop a permutation algorithm that can work on the EQF elements or on the QTL intervals to compute a range of EQF thresholds that vary from strict to liberal for assessing the significance of QTL hotspots. For better understanding, we use a simple example in Figure 1 to illustrate the scheme of our statistical procedure from using the QTL intervals (matrix) to obtaining the permutation EQF thresholds. Real example and simulation study are followed to demonstrate the capability of our procedure, investigate the detection properties, and also compare with the methods using individual-level data in QTL hotspot detection.

**Figure 1.**
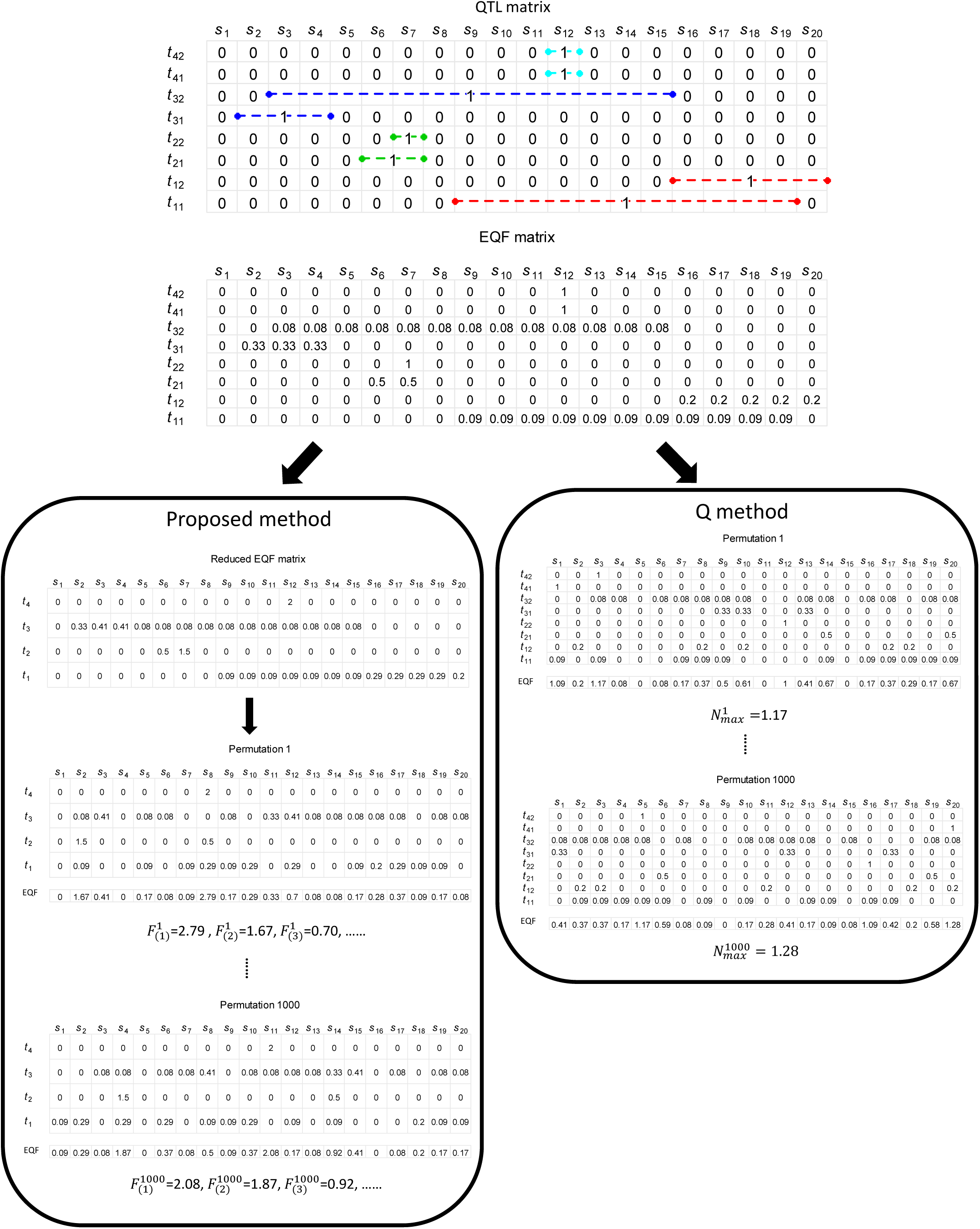
The schemes of the proposed statistical procedure and the Q-method in obtaining the permutation thresholds. The QTL data from the public database are first recorded into a QTL matrix, where the QTL intervals take a value of one and the remaining elements will be treated as zeros at the corresponding positions. The output of the analysis is a QTL matrix, where columns represent the genomic positions (*s*_i_’s) and rows represent the traits (*t*_ij_ denotes the *j*th trait of the *i*th trait category). Then, the QTL matrix is converted into the expected QTL frequency (EQF) matrix by using the uniform distribution method. The Q-method permutes the EQF matrix for each trait (the column cells) to obtain the permutation EQF threshold, *β* (see text). The proposed procedure groups together the related traits into the same trait categories and pools their EQF values to form a reduced EQF matrix, and the reduced EQF matrix for each trait category (the column cells) are permuted to obtain a series of EQF thresholds, *γ*_n,a_’s (see text).

### The QTL matrix

The QTL intervals contain complete information about the QTLs that need for hotspot detection. We use a row array of the same length as the genome size to summarize the QTL intervals for each trait. For each trait, we regularize the QTL intervals into the elements of a row array as follows: Each QTL interval corresponds to an element of the length as its width at the corresponding position, and a value of one is assigned to the element. The remaining elements will be treated as zeros. In this way, the elements in the row array are either one or zero with unequal lengths (see Figure 1 for graphical illustration). Combining the arrays for all traits will form a QTL matrix with different element sizes. The QTL matrix (an atypical matrix) will be used to construct the EQF matrix for permutation as described below.

### The expected QTL frequency

Consider that a QTL matrix has been constructed from the database. We assume that there are *T* traits mapped for *N*_1_, *N*_2_, …, *N*_*T*_ QTLs, respectively, in the experiments (*N*_QTL_ = *N*_1_ + *N*_2_ + … + *N*_*T*_), and that the genome is divided into *S* sequential equally spaced bins, each with size Δ (say Δ=0.5 cM), for investigation. We use the uniform distribution to compute the EQF value of each bin over the total experiments for hotspot detection. For a bin (*x*, *x* +Δ) and a QTL interval (*a*, *b*), where *x*, *a* and *b* denote the genome positions, there are two possible relationships between them: (1) they have an overlap, i.e. (*a*, *b*) ∩ (*x*, *x* +Δ) ≠ ∅, or (2) they have no overlap, i.e. (*a*, *b*) ∩ (*x*, *x* +Δ) = ∅. Only the QTL intervals having overlaps with a bin contribute to the EQF value of this bin, and such a QTL is called a contributive QTL of a bin. By assuming that the QTL position is uniformly distributed over its own QTL interval, the probability of a contributive QTL localized in a bin is the ratio between the lengths of the overlap and the interval. Summing over the probabilities of all the contributive QTLs in a bin will give the EQF value of this bin over all traits. Explicitly, we define *F*_*s*_ as the EQF value of the *s*th (*s* = 1, 2, …, *S*) bin between *x* and *x* +Δ for all traits over all experiments by

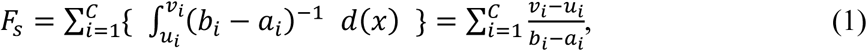

where *C* is the number of the contributive QTLs of the bin (*x*, *x* +Δ), (*u*_*i*_, *v*_*i*_) is the overlap region, (*a*_*i*_, *b*_*i*_) denotes the *i*th QTL interval, and (*b*_*i*_ − *a*_*i*_)^−1^ is the uniform distribution density function over the interval. A ratio (*v*_*i*_ − *u*_*i*_)^⁄^(*b*_*i*_ − *a*_*i*_) will be added to the EQF of the bin despite of the effect size of a contributive QTL. If a QTL interval fully covers the bin, (*v*_*i*_ − *u*_*i*_) =Δ and the ratio (probability) is Δ/(*b*_*i*_ − *a*_*i*_). If a QTL interval is fully covered by the bin, (*v*_*i*_ − *u*_*i*_) = (*b*_*i*_ − *a*_*i*_) and the probability is 1.

The computation of the EQF values in equation (1) can be illustrated by Figure 1 and the example in Figure 2. In Figure 2, we use a set of selected 196 QTLs (*N*_QTL_ = 196) in the 1^st^ chromosome from GRAMENE databases for illustration. These 196 QTLs are responsible for 95 different traits belonging to four of the nine trait categories. Depending on the bin position, the number *C* involved in computing *F*_*s*_ is different for different bins. For example, *C* = 4 for a bin between 80~90 cM as the region is only covered by the four same QTL intervals, and *C* = 22~63 for a bin between the 140~160 cM as the within bins may overlap with different QTL intervals. By equation (1), the EQF value of a bin in the 80~90-cM region is 0.037, and the EQF value of a bin in the 140~160-cM region is 0.91~7.38. The EQF value can be calculated at every bin to produce an EQF hotspot architecture along the chromosomes as shown in Figures 2 and 3. A higher EQF value reflects a greater expectation of localizing a QTL in a bin. A hotspot detection is claimed in a bin if its EQF value is higher than a specified threshold that will be determined by permutation tests as given below.

**Figure 2.**
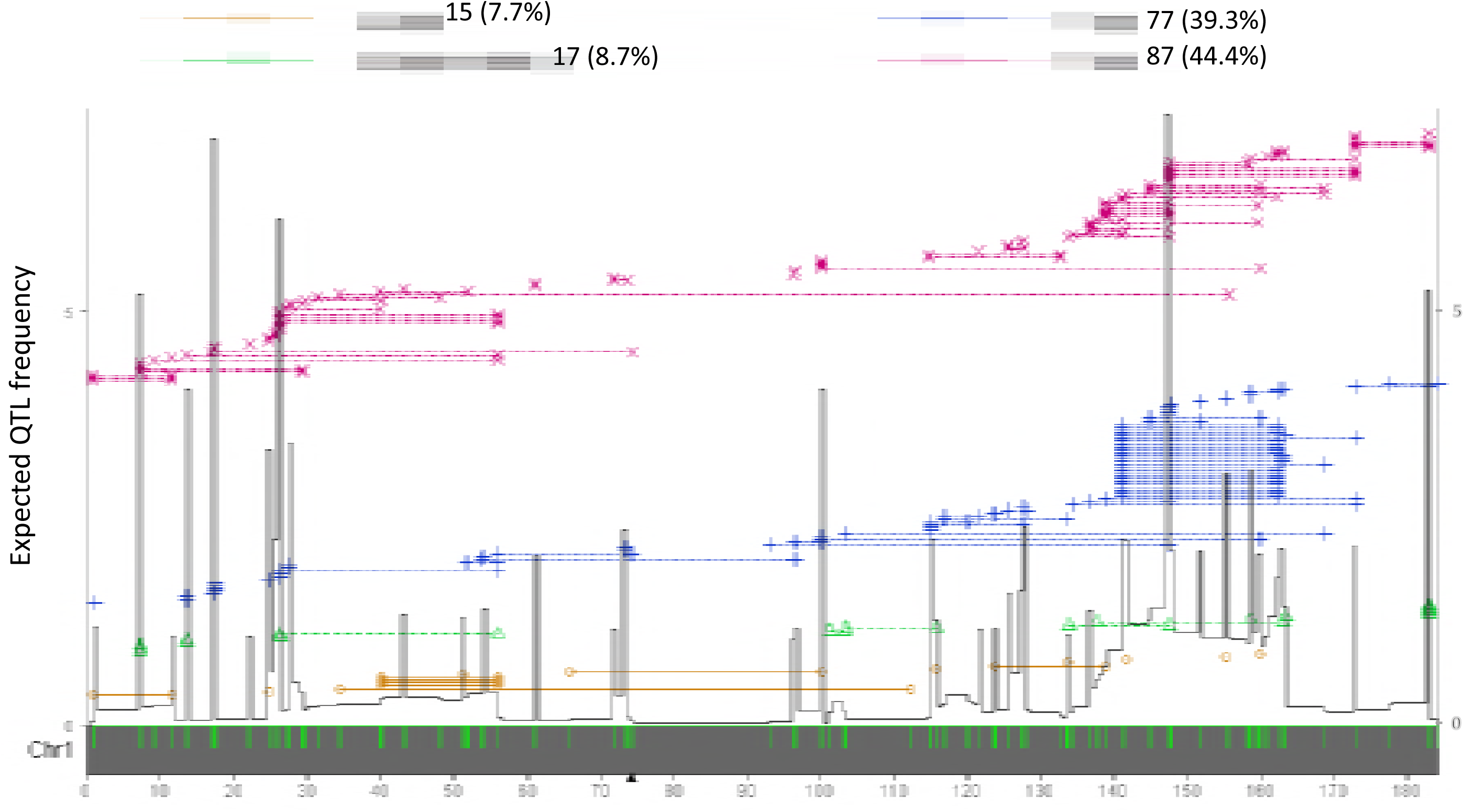
An illustration of the QTL data structure and the uniform method of computing the expected QTL frequency (EQF) in hotspot detection. The 196 QTLs in the rice 1^st^ chromosome from Gramene Rice database (http://www.gramene.org/) are used for illustration. The green ticks on the x-axis denote the positions of the 163 markers. The dotted lines denote the lengths of the marker intervals flanking the QTLs responsible for yield, vigor, sterility and quality traits (denoted as x, +, ∆ and ○, respectively). The EQF architecture (the black line) are constructed by the uniform method with bin size of 0.5 cM. The black lines on the right and left borders represent the EQF. ▲denotes the position of centromere.

**Figure 3.**
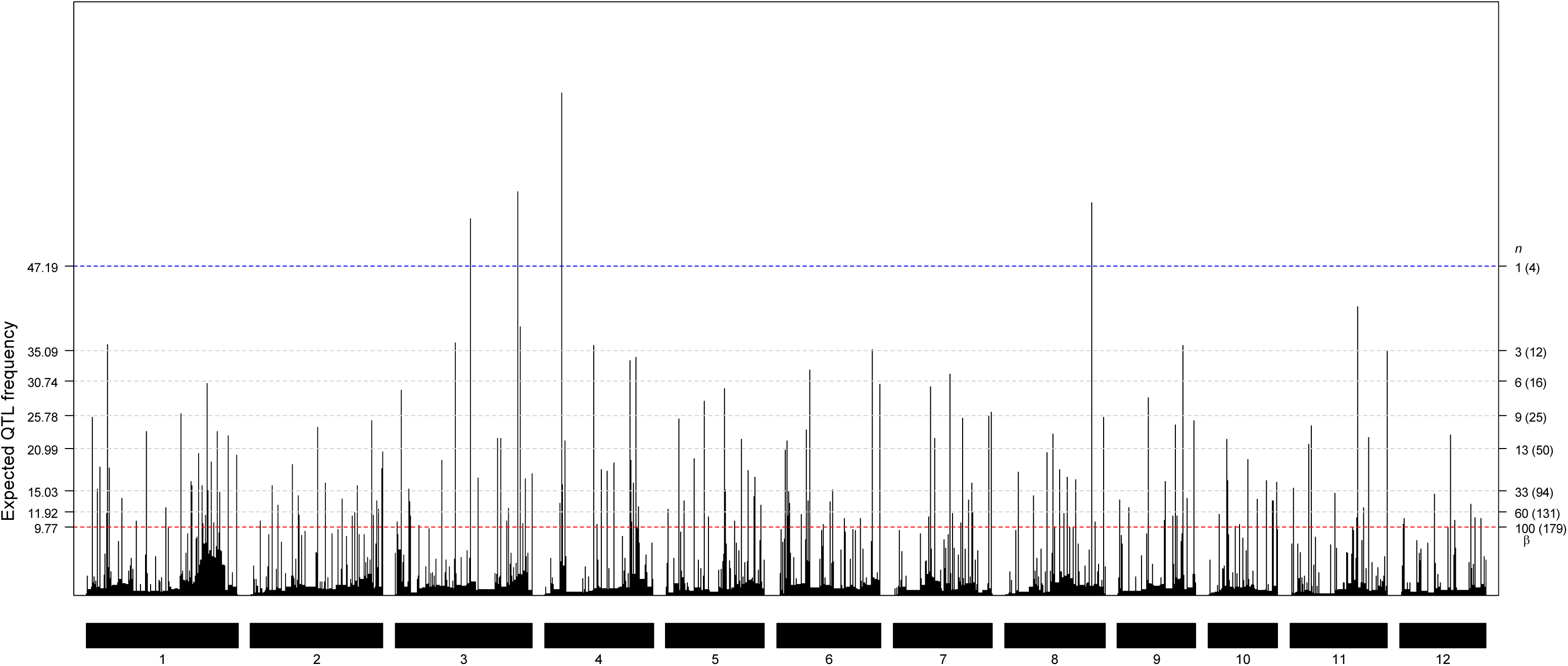
The EQF architectures along the 12 chromosomes and the hotspots detected under different EQF thresholds (*γ*_n,0.05_) associated with their qFreq(*n*) statistics at GWER of 5%. The thresholds *γ*_n,0.05_ are coordinately represented by the left and right axes. The left axis denotes the values of EQF, and the right axis denotes the values of *n*. The blue line corresponds to the EQF threshold *γ*_l,0.05_ = 47.19 for the qFreq(1) statistic of detecting at least one hotspot, and there four significant hotspots with *γ*_l,0.05_. The red line shows *γ*_l00,0.05_ = 9.77 for the qFreq(100) statistic of detecting at least 100 hotspots, which approximately corresponds to *β* (the EQF threshold of the Q-method), and there are 179 significant hotspots with *γ*_l00,0.05_.

### The EQF matrix

Equations (1) is to compute the EQF value of a bin for all traits over all experiments in the genome. It is also desirable to compute the EQF value of a bin for each single trait, and further to construct the EQF matrix for permutation to determine the thresholds. Let *f*_*ts*_ denote the EQF value of the *s*th bin for the *t*th trait. It is straightforward to obtain *f*_*ts*_ using equation (1) by simply replacing the number of contributive QTLs for all traits, *C*, by the number of contributive QTLs for the *t*th trait,

*C*_*t*_. That is

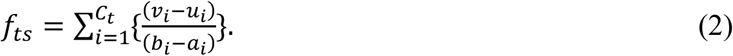

We now define the EQF matrix as ***F*** = {*f*_*ts*_}_*T*×*S*_, where *t* = 1, 2, …, *T* is the index for row dimension, and *s* = 1, 2, …, *S* is the index for column dimension. We have 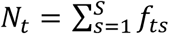 is the number of QTLs for the *t*th trait, and 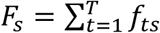 is the EQF value of the *s*th bin for all traits. By pooling the EQF values of genetically correlated traits, the ***F*** matrix will be further converted into a reduced ***F*** matrix (row dimension less than *T*) for permutation to determine the EQF thresholds for assessing the significance of hotspots.

The current methods of West *et al.* (2007), Wu *et al.* (2008) and Li *et al.* (2010) permute the QTL matrix for hotspot detection in the genetical genomics study. Their methods can be directly implemented to permute the EQF matrix for QTL hotspot detection using public databases. We follow Neto *et al.* (2012) to still call these methods implemented to the EQF matrix “Q-method”. The Q-method will permute the row elements of the EQF matrix separately by traits and then obtain the EQF sums over all traits for every bin in the genome, and the maxima of EQF sums in all permutations are used to compute the EQF threshold for assessing the QTL hotspots (Figure 1). Similarly, the Q-method does not account for the correlation structure among traits, and hence it will severely underestimate the null distribution of hotspot sizes (thresholds) and detect too many spurious hotspots (Neto *et al.* 2012; also see below). To consider the correlation structure among traits, we group together the correlated traits into the same trait categories and pool together their EQF values to form a reduced EQF matrix for permutation. For example, the panicle numbers, grains per panicle and grain weight are all related to yield production in rice, and their EQF values can be summed and become the EQF value of the yield trait category to form a reduced EQF matrix (the row dimension reduced by 2 to become *T–*2). As pleiotropy and linkage of genes are the genetic causes of correlations between traits (Falconer and Mackay 1996), the trait grouping intends to pool and permute these QTLs together, so that to cope with the underestimation of threshold. Other grouping strategies can be also considered (see CONCLUSION AND DISCUSSION). Permutation with trait grouping can effectively obtain much stricter thresholds to prevent detecting spurious hotspots, as will be shown below. Furthermore, inspired by the work of Neto *et al.* (2012), rather than examine the possible spurious hotspot sizes, we consider the possible spurious hotspot numbers in the genome to present a permutation algorithm for computing the thresholds. Our permutation algorithm shuffles the reduced EQF matrix to compute a series of EQF thresholds ranging from strict to liberal for assessing the QTL hotspots.

### The permutation algorithm

For a fixed hotspot number *n* = 1, …, *k,* with *k* the hotspot number delivered by the Q-method (see below), across the genome, we first define qFreq(*n*) as the *n*th EQF sum of the *S* ordered observed EQF sums (namely: *F*_(1)_, *F*_(2)_, …, *F*_(*s*)_, ordered from highest to lowest) in the reduced ***F*** matrix and use it as a test statistic for at least *n* spurious hotspots under the null hypothesis that the QTLs are randomly distributed in the genome. We describe the permutation algorithm that can control the GWER of detecting at least *n* hotspots at a fixed α level as follows:

1. For each trait (category), the EQF values of the *S* locations (bins) in the reduced ***F*** matrix (with row dimension ≥ 2) are permuted to generate a new permuted EQF matrix ***F***∗. That is, the elements in each row of the reduced ***F*** matrix are swapped to produce the ***F***∗ matrix (see Figure 1). As every row element can be only assigned to one of the *S* bins, the row sums of the permuted and observed matrices are the same and are equivalent to *N*_*t*_ for the *t*th trait (category), *i.e.*,

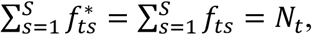

where 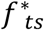 is the element of ***F***^∗^.
2. Compute the total EQF sums over all traits (categories) for the *S* locations, *i.e*. 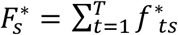, and order them 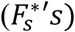 from highest to lowest as 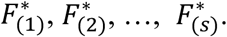.
3. For a fixed hotspot number *n*, obtain and store 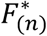 corresponding to the *n*th EQF sum of the *S* ordered EQF sums for ***F***^∗^.
4. Repeat steps 1–3 *B* times so that there are *B* new permuted matrices (namely, ***F***^1^, ***F***^2^,…, ***F***^*B*^) for obtaining the associated 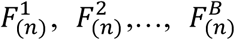. The *B*-permutation samples of 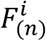, is an estimate of the null distribution of the test statistic qFreq(*n*) for at least *n* spurious hotspots anywhere in the genome, given that the QTLs are randomly distributed along the genome.
5. The upper (1-α)-quantile of the *B*-permutation samples generated in step 4 is the EQF threshold, denoted by *γ*_*n*,*α*_, of the test statistic qFreq(*n*) for assessing QTL hotspots.

The EQF threshold *γ*_*n*,*α*_ can control GWER of qFreq(*n*) at level α for detecting at least *n* spurious hotspots somewhere in the genome under the null. That is

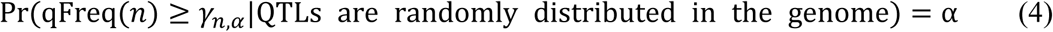

The quantity α is the probability of detecting exactly *n* spurious hotspots somewhere in the genome under the null. However, similar to the argument in Neto *et al.* (2012), the threshold *γ*_*n*,*α*_ can also control the full-null GWER for more than *n* hotspots, as detecting more than *n* hotspots is less likely than detecting *n* hotspots given the threshold *γ*_*n*,*α*_. Therefore, by adopting *γ*_*n*,*α*_ we can control GWER of qFreq(*n*) at level α of detecting at least *n* spurious hotspots under the null.

### Circular genomic permutation

The basic idea of the above algorithm is to randomly shift the EQF bins along the genome to obtain the thresholds for QTL hotspot detection. When doing this, a QTL interval will be broken into several bins for permutation. To keep the QTL intervals intact in permutation, we can consider the genome to be circular (Cabrera *et al.* 2012) and directly implement the proposed algorithm to randomly swap the QTL intervals of correlated traits together in the circular genome. Equation (1) is then used to compute the EQF sums of all bins in each permutation for obtaining the EQF thresholds. We call this circular permutation framework the QTL-interval permutation in our statistical procedure. In this way, the proposed algorithm can deploy both the EQF-bin permutation and the QTL-interval permutation to compute a series of thresholds, *γ*_*n*,*α*_’s, for qFreq(*n*)’s to assess the significance of QTL hotspots.

In general, performing permutation on the EQF bins or the QTL intervals for the hotspot detection is not computationally demanding at all. On the contrary, performing permutation on individual-level data, such as in the approaches of the Breitling *et al.* (2008) and Neto *et al.* (2012), is computationally expensive and requires parallel computations on a cluster (Neto *et al.* 2012), mainly because it involves repeated QTL mapping analysis for thousands of traits in each permutation. Briefly, the proposed statistical procedure for genome-wide hotspot detection includes four steps: (1) obtaining the QTL intervals (matrix) from the database; (2) constructing the EQF matrix from the QTL matrix using the uniform distribution method; (3) building the reduced EQF matrix by grouping of correlated traits into the same trait categories; (4) computing the thresholds by implementing the permutation algorithm on the reduced EQF matrix or on the QTL intervals (with trait grouping). Our statistical procedure actually generalizes and combines the ideas of the previous works on QTL hotspot detection from using genetical genomics experiments to using public databases. The value of *n* is allowed to vary from 1 to *k*. Given *β* as the threshold value of the Q-method, *k* is determined by *β* = *γ*_*k*,*α*_. The *k* thresholds, *γ*_1,*α*_, *γ*_2,*α*_,…, *γ*_*k*,*α*_, range from the most conservative to the most liberal. Then, our statistical procedure of using *γ*_*k*,*α*_ as the threshold is equivalent to the Q-method deployed for permuting the original EQF matrix, which will also deliver the most liberal EQF threshold and suffer from excessive spurious hotspots. With the grouping strategy and permutation algorithm, our statistical procedure can provide a range of more conservative thresholds to prevent the detection of spurious QTL hotspots as will be validated in the next section.

## REAL EXAMPLE ANALYSIS AND SIMULATION STUDY

In this section, real example with comparative analysis and simulation study are conducted to illustrate the proposed statistical procedure, investigate the property of the proposed procedure, evaluate the performance of the proposed procedure, and compare with other methods in QTL hotspot detection. In real example analysis, the QTL data collected in GRAMENE rice database were analyzed to detect QTL hotspots, and then a genome-wide comparative analysis of the detected QTL hotspots and the known genes collected in Q-TARO rice database was performed for cross validation and practical use. In simulation study, we investigate the GWERs of the proposed statistical procedure and the Q-method under different levels of correlation among traits, and assess their performances in the detection of QTL hotspots. The proposed statistical procedure operates on summarized QTL data instead of original individual-level data for hotspot detection. There must be some information loss between the two types of data during the detection process. We implement our procedure, Q-method, Breitling’s method (2008) and Neto’s approach (2012) to analyze a simulated genetical genomics data set, and compare their differences for examining such information loss in hotspot detection.

### The GRAMENE rice database example

#### The GRAMENE database and Q-TARO database

Both the GRAMENE database and Q-TARO database are web-accessible and common reference databases for rice research. The GRAMENE database collects 8216 QTLs (*N*=8216) responsible for 236 different traits (*T*=236) from 230 published studies (experiments). These 236 traits are further classified (grouped) into the nine trait categories (*T*=9) according to the general agronomic consideration (see Tables S1 and S2 in supplementary material). The nine trait categories including yield, vigor, anatomy, development, abiotic stress, quality, sterility or fertility, biotic stress, and biochemical traits. For example, the yield category includes traits, such as the panicle numbers, grains per panicle and grain weight, etc. The nine trait categories include 26, 15, 47, 12, 52, 44, 11, 8 and 21 component traits, respectively, and they contain 1956, 1767, 1267, 901, 767, 555, 470, 378 and 155 QTLs, respectively. The total length of the rice 12 chromosomes is ~1536.9 cM (Harushima *et al.* 1998; Matsumoto *et al.* 2005). The 1^st^ chromosome has the highest QTL density with ~6.77 QTLs per cM, and the 12^th^ chromosome has the lowest QTL density with ~3.67 QTLs per cM. The QTL density is ~5.35 QTLs per cM across the 12 chromosomes.

The flanking marker pairs of the 8216 QTLs (8216 QTL intervals) are recorded (see the S1_DataQTL file in the supplementary material) and used for detecting QTL hotspots. The detection results will be investigated and compared with the 122 known genes (see the S2_DataGene(NV) file in supplementary material) identified through natural variation methods (such as using the experiments of rice cultivars, landraces, or wild relatives) in Q-TARO database (Yamamoto *et al.* 2012). We first projected the two physical maps of the GRAMENE rice and Q-TARO databases into a consensus genetic map by homothetic function (Chardon *et al.* 2004) with their common markers (Harushima *et al.* 1998; Matsumoto *et al.* 2005). There are 1914 common markers (see the S1_DataQTL file) on the consensus map with an average marker density of one marker every 0.81 cM. By using equations (1) with bin size of 0.5 cM (Δ=0.5 cM), the EQF architectures (Figure 3) and the EQF matrix can be obtained. The EQF matrix has a dimension 236×3070, and the reduced EQF matrix has a dimension 9×3070 after grouping of the 236 traits into the nine trait categories. Both the EQF-bin permutation and QTL-interval permutation are considered in the analysis. The detection results are summarized and presented below and in the supplementary material.

#### QTL hotspots

The thresholds *γ*_*n*,0.05_, *n*=1, 2,…, *k*, obtained by permuting the EQF bins and the QTL intervals are very similar (see Figure S1). Figure 3 presents the EQF architectures of the 12 chromosomes and the hotspots detected under different EQF thresholds (*γ*_*n*,0.05_) obtained using the EQF-bin permutation. In Figure 3, the threshold values *γ*_*n*,0.05_’s for the test statistic qFreq(*n*)’s are coordinately represented by the left and right axes. For example, the 1^st^ highest peak (on the 4^th^ chromosome) has an EQF value 71.97, and therefore qFreq(1) for detecting at least one hotspot in the genome is 71.97 (qFreq(1) =71.97). The threshold for qFreq(1) is *γ*_1,0.05_, which is 47.67 (*γ*_1,0.05_ = 47.67). Since qFreq(1) > *γ*_1,0.05_, it means that there exists at least one hotspot in the genome, and in practice there are four significant hotspots (on the 3^rd^, 4^th^ and 8^th^ chromosomes). Similarly, qFreq(3) = 56.30 (the 3^rd^ highest peak on the 8^th^ chromosome) is the statistic for detecting at least three hotspots, and *γ*_3,0.05_ =35.29 is the threshold for qFreq(3). As qFreq(3) > *γ*_3,0.05_, it means that there are at least three hotspots in the genome, and in practice there are ten significant hotspots (on the 1^st^, 3^rd^, 4^th^, 6^th^, 8^th^, 9^th^ and 11^th^ chromosomes). The highest peak of the 1^st^ chromosome has an EQF value 35.89 and was significant under *γ*_3,0.05_ =35.29, but not significant under *γ*_1,0.05_ =46.70. The EQF value of the 6^th^ highest peak (on the 3^rd^ chromosome) is 38.52, *i.e.* qFreq(6)=38.52, and *γ*_6,0.05_ is 30.66. Since qFreq(6) > *γ*_6,0.05_, it means that there are at least six hotspots in the genome, and in practice there are 16 hotspots detected. Likewise, qFreq(9) > *γ*_9,0.05_, meaning that there are at least nine hotspots in the genome, and in practice there are 25 significant hotspots. The top 100 hotspots are significant under *γ*_39,0.05_ =14.21. Chardon *et al.* (2004) empirically suggested 5 times of the average EQF value per bin (5.35÷2× 5 = 13.38) as the threshold, which roughly corresponds to *γ*_46,0.05_ =13.29, and there are 116 significant hotspots detected under *γ*_46,0.05_. The EQF threshold obtained by the Q-method is about 9.77 (corresponding to *γ*_100,0.05_, where 100 is the upper bound of *n*, *i.e. k*=100), leading to the detection of 179 QTL hotspots, among which many of them are believed to be spurious since the correlations among traits are not considered by the Q-method.

#### Effects of bin sizes and QTL interval sizes

The above analyses use a bin size of 0.5 cM (Chardon *et al.*, 2004) and all the 8216 QTL intervals for QTL hotspot detection. It is of interest to assess the effects of bin size and removing large QTL intervals on the hotspot detection. The thresholds of *γ*_*n*,0.05_, *n*=1, 2,…, 100, by using bin size of 1 cM and removing the QTL intervals larger than 10 cM (10+ cM intervals) in the analyses are shown in Figure S1. Using a bin size of 1 cM, it is found that the thresholds are higher, and the EQF architectures show similar profiles to those of using a bin size of 0.5 cM with higher EQF values at the same or very nearly the same positions (not shown). For example, using a bin size of 1 cM (0.5 cM) with all intervals, the thresholds *γ*_1,0.05_ and *γ*_60,0.05_ are 53.77 (47.67) and 15.36 (11.89), and there are 4 (4) and 124 (131) detected hotspots. Under *γ*_1,0.05_, the same four hotspots are detected for the both bin sizes. Under *γ*_60,0.05_, most of the detected hotspots are the same. Other bin sizes, such as >1 cM, can be also considered. We suggest to use a bin size about equal to the marker density for hotspot detection (the average marker density is 0.81 cM in our case). In general, as bin sizes increase, the hotspot resolution will decrease, and the EQF values for every bin and the permutation thresholds will increase.

The medium, mean and SD of the interval sizes are 0.56, 9.82 and 16.82 cM, respectively, showing that the QTL intervals vary greatly in size. Among the 8216 QTLs collected in GRAMENE database, 309 (3.76%) QTLs are localized at markers, 3791 (46.14%) QTLs are localized in the marker intervals with sizes between 0 and 0.5 cM, 274 (3.33%) QTLs are localized in the 0.5-2 cM intervals, 509 (6.20%) QTLs are in the 2-5 cM intervals, 744 (9.06%) QTLs are in the 5-10 cM intervals, and 1023 (12.45%) QTLs are in the 10-20 cM intervals. Therefore, there are 2589 (about 31.15%) QTLs with interval sizes >10 cM. Researchers may only consider those QTL intervals with small sizes, say ≦10 cM, in the analysis. If the 10+ cM QTL intervals are not considered in the analysis, the EQF values at very bin are smaller (not shown) and the permutation thresholds decrease slightly (see Figure S1), mainly because the 10+ cM QTL intervals contribute For example, if the 10+ cM QTL intervals are excluded, *γ*_1,0.05_ and *γ*_60,0.05_ become 46.14 and 11.05, respectively, and there are 4 and 132 hotspots detected. Under *γ*_1,0.05_, the four detected hotspots are the same. Under *γ*_60,0.05_, an extra hotspot is detected by removing the 10+ QTL intervals from the analysis.

For all thresholds *γ*_*n*,0.05_, *n* = 1, 2, …, *k*, the observed hotspot numbers are found to exceed the expected hotspot numbers, very likely because the traits in different categories are still correlated to some extent after grouping (see CONCLUSION AND DISCUSSION). The proposed statistical procedure allows to use the *k* statistics associated with their respective thresholds ranging from high to low for broad consideration in assessing the significance of QTL hotspots. The results indicate that there are many significant hotspots in the GRAMENE rice database. It would be interesting to further explore the genes underlying these hotspots. We then perform a genome-wide comparative analysis of the detected QTL hotspots and the 122 known genes in Q-TARO databases about their locations and functions.

#### QTL hotspots, gene locations and gene functions

The comparative analysis reveals that many of the detected hotspots are localized in the vicinity of the genes for their associated traits (see Figures S2). Taking the top 131 hotspots significant under *γ*_60,0.05_ =11.89 (obtained by the EQF-bin permutation) as an example. Among the 122 genes, 17 genes overlap with the hotspots (the distances between them are zeros), 48 genes are localized within a distance of 1 cM or less from their nearest hotspots with an average distance of 0.23 cM (standard deviation 0.28 cM), 78 genes are localized within a distance of 2.5 cM or less from the their nearest hotspots with an average distance of 0.83 cM (SD 0.84 cM), and 22 genes have distances of 2.5 to 5 cM to their closest hotspots with an average distance of 3.31 cM (SD 0.68 cM). To perform a statistical test for the conformity in locations between genes and hotspots, we assume that the number of overlap, *y*, follows hypergeometric distribution (*N,r,n*)

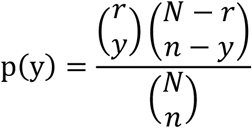

(Peng *et al.* 2010; Cabrera *et al.* 2012), where *N* is the number of bins, *n* is the number of detected hotspots and *r* is the number of known genes. In this case, the *N* = 3070, *n* = 131, *r* = 102 and *y* = 17. Note that *r* is assigned to the number of 102 instead of 122 because of gene overlaps. For example, six genes Pik-p, Pik-1, Pik-2, Pik-m, Xa3 and Xa26 overlap at the end of the 11^th^ chromosome (see Figure S2). The p-value is p(y ≥ 17)=1.497 × 10^−7^, which indicates that QTL hotspots have the tendency to co-localize with genes (known genes and detected hotspots are conformable in location). If the QTL-interval permutation is used, the value of *γ*_60,0.05_ is 12.08, and there are 129 QTL hotspots detected. Two less QTL hotspots are detected due to the use of higher threshold, and the p-value is more extreme.

These genes are also functionally related to the QTL components in the hotspots. For example, the *PSR* gene responsible for shoot regeneration is located in the hotspot [1,73-73.5] (at 73~73.5 cM of the 1^st^ chromosome) with EQF value 23.47 for all traits and EQF value 11.14 for vigor traits. The *dth3* gene for days to heading is located in the hotspot [3,6.5-7] with EQF value 29.41 for all traits and EQF value 13.92 for development traits. The *Hd16* gene acting as an inhibitor in the rice flowering pathway is located in the hotspot [3,152-152.5] with EQF value 38.52 for all traits and EQF value 28.23 for sterility traits. The *WFP* gene for regulating panicle branching and grain yield is located very close the hotspot [8,105.5-106] with EQF value 56.30 for all traits and EQF value 29.18 for vigor traits. The *GPS* gene for leaf photosynthesis rate and yields is located in the hotspot [4,110.5-111] with EQF value 34.13 for all traits, EQF value 6.83 for vigor traits and EQF value 7.11 for yield traits. The well-known pleiotropic gene *SCM2* responsible for lodging resistance and yield is located very close to the detected hotspot [6,115.5-116], which is mostly composed of the QTLs for yield, vigor and development traits. The *Xa3* gene conferring resistance to bacteria blight is located at the hotspot [11,118-118.5] with EQF value 35.11 for all traits, EQF value 13.00 for development traits and EQF value 12.04 for vigor traits. The analysis also recognizes several hotspot regions that do not have any known gene nearby so far (for example, in the 40~110-cM region of the 5^th^ chromosome). These regions can be considered as potential regions of new genes that are functionally related to the QTLs in the hotspots. In general, the QTLs in the hotspots and their nearby genes are related in functions and locations to their associated traits.

### Simulation study

The simulation study includes three parts. The first part aims to assess and compare the GWERs of the proposed statistical procedure and the Q-method in hotspot detection using the QTL data with correlation structure under the null hypothesis of no QTL hotspots. The second part focuses on assessing the performances of the two methods in detecting QTL hotspots when QTL hotspots are present. These two parts consider a 100-cM chromosome. The ~1900 marker intervals and the 8216 QTL intervals in GRAMENE rice database (see the S1_DataQTL file) are served as the sample populations to generate the markers and QTLs onto the simulated chromosomes. The third part attempts to evaluate the information loss when using summarized data instead of individual-level data by comparing the performances of the different methods in analyzing a simulated genetical genomics data set. The bin size in detection analysis is 0.5 cM (Δ=0.5) for the first two parts and is 2 cM (Δ=2.0) for the third part.

#### Genome-wide error rates

In the first part, the QTLs are generated to be randomly distributed but with different levels of correlation. We first assume the QTLs belong to two trait categories and each trait category contains 150 QTLs for the two different traits. Then, we deploy a hierarchical two-stage process to generate the QTL data in a trait category: (1) 105 QTLs are randomly placed to the 200 possible positions (bins) without coincidence to determine their positions; (2) The remaining 45 QTLs are then randomly assigned to the 105 determined QTL positions (in the first stage) by allowing coincidence. The hierarchical two-stage process can generate randomly distributed QTLs (traits) with correlation structure in the genome. The first stage is to generate randomness for QTLs, and the second stage allows to create correlations among QTLs. We denote the above process as a (105,45) process for generating the QTLs in a trait category. Then, the QTL data with different strengths of correlation under the null can be generated using different process, such as the (60,90) or (150,0) process. It follows that the QTLs generated by the (60,90) process will be more correlated as compared to those by the (105,45) process, and those generated by the (150,0) process are uncorrelated under the null. We consider four kinds of processes, (A) (150,0) and (150,0), (B) (135,15) and (105,45), (C) (105,45) and (60,90), (D) (60,90) and (15,135), to generate the 300 QTLs in the two trait categories. After the QTL positions are determined, the lengths of their flanking markers are randomly sampled from the 8216 QTL intervals in the GRAMENE database. Under such settings, the QTLs in the different trait categories are uncorrelated, and those in the same trait category have different strengths of correlation. The four QTL data sets from the processes (A), (B), (C) and (D) are uncorrelated, weakly correlated, moderately correlated and highly correlated, respectively, and the data sets are analyzed by the proposed statistical procedure (*T*=2) and Q-method (*T*=4) to assess their GWERs. Both the EQF-bin and QTL-interval permutations are considered in the analysis. In our statistical procedure, we set *k*=10 to investigate the GWERs of qFreq(*n*), *n* = 1,2, …,10, here. The number of simulated replicates is 1000.

The EQF-bin and QTL-interval permutations produce the similar permutation thresholds and same results. The results of the EQF-bin permutation are presented here. Figure 4 displays the observed GWERs of the proposed procedure and the Q-method at the α =0.02, 0.04, 0.06, 0.08 and 0.10 levels for the four QTL data sets. Figures 4A shows that for uncorrelated QTLs the observed GWERs of the two approaches are about the right target levels. When the QTLs are correlated, Figures 4, B-D show that the GWERs of the Q-method are higher than the target levels and increase with correlation strength. Most strikingly, for the highly correlated data, the observed GWERs of the Q-method are closed to 1 at all the levels (Figure 4D), which implies that the detected hotspots by the Q-method are very likely to be spurious without accounting for the correlation features. On the other hand, the proposed procedure can cope with the correlations among QTLs (by trait grouping) and control the GWERs close to the target levels.

**Figure 4.**
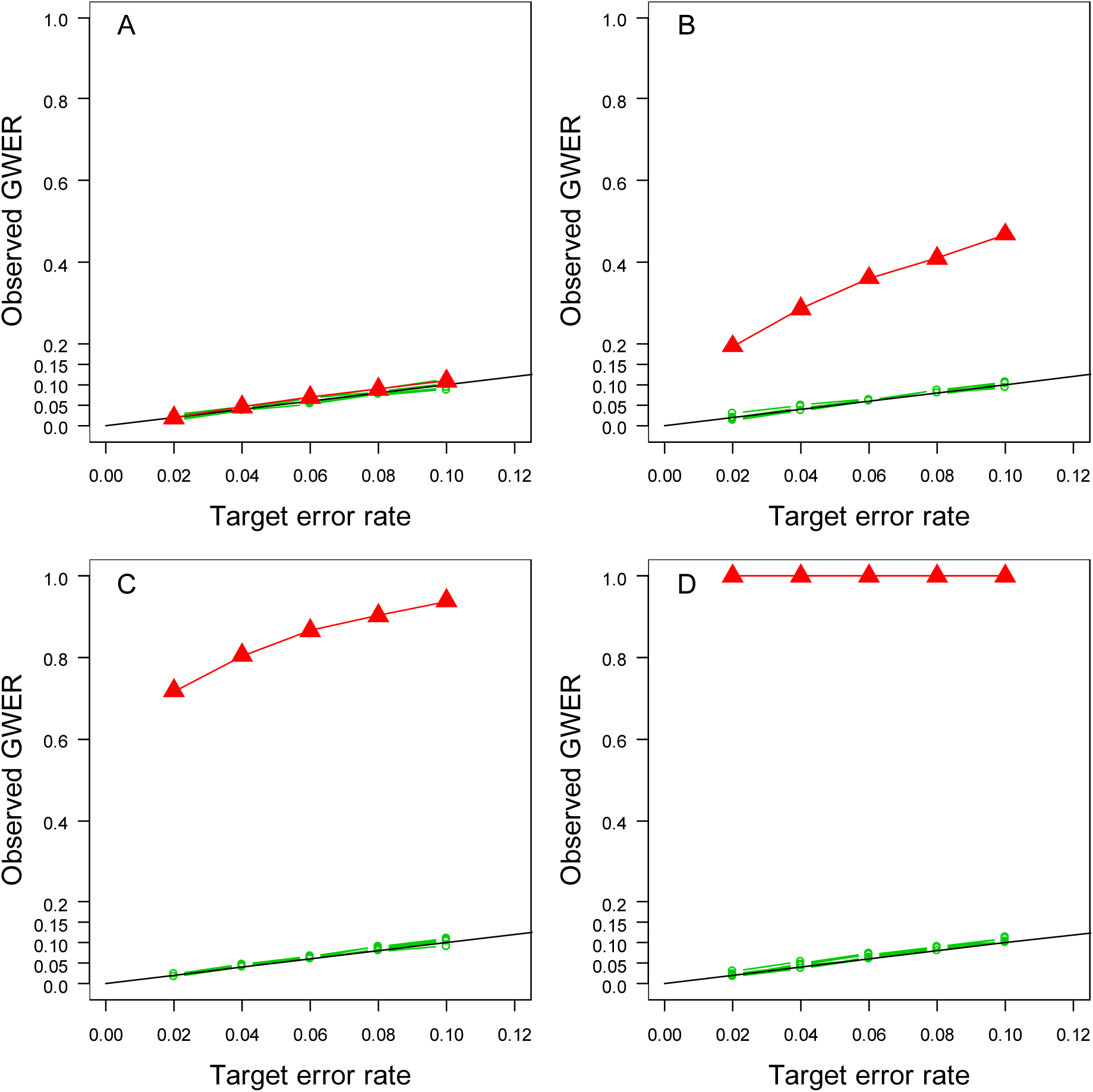
Observed GWERs for the qFreq(*n*), *n* = 1,2, …, 10, statistics of the proposed statistical procedure and for the Q-method at the α =0.02, 0.04, 0.06, 0.08 and 0.10 levels under varying strengths of QTL correlation: (A) uncorrelated, (B) weakly correlated, (C) moderately correlated, and (D) highly correlated, respectively. Black lines show the targeted error rates. Red lines show the observed GWERs of the Q-method. Green lines show the observed GWERs of the qFreq(*n*), *n* = 1,2, …, 10, statistics.

#### Performance in QTL hotspot detection

For the second part, we assume there are six hotspots in the same 100-cM chromosome, and each hotspot is caused by a single gene. The six hotspots are assumed to be located at 10.25, 20.35, 31.15, 47.25, 56.40, and 67.20 cM of the chromosome, respectively. To generate QTLs onto the chromosome, the contributive QTLs of the top 100 hotspots in GRAMENE database are served as the sample populations of the six hotspots. The number of contributive QTLs, *C*, ranges from 29 to 330 in the top 100 hotspots. For each simulated replicate, the contributive QTLs in the top 100, top 20, top 50 to 100, top 30 to 60, top 50, and top 10 hotspots are served as the sample populations of the 1^st^, 2^nd^, 3^rd^, 4^th^, 5^th^ and 6^th^ hotspots, respectively. Such settings imply that the 2^nd^ and 6^th^ hotspots are stronger hotspots, and the 3^rd^ hotspot is a weaker hotspot. Once the number *C* is determined, the relative positions and interval lengths of the *C* QTLs are then randomly chosen from the respective sample population. The trait names and categories of the sampled QTLs are still used, and hence correlation structure between QTLs is similar to the GRAMENE database. The number of simulated replicate is 1000. For each simulated data set, the EQF matrix (*T*≈236) and reduced EQF matrix (*T*≈9) are constructed for detecting the QTL hotspots.

To assess the performances of the two methods in detecting QTL hotspots, we use the true positive rate (TPR) and positive predictive value (PPV) to jointly measure the correct detection rate, and use the false discovery rate (FDR) to measure the incorrect detection rate. The TPR defines the proportion of the six hotspots that are correctly detected, and the PPV (FDR) defines the proportion of true (false) positive detection among all positive detections in the chromosome over the 1000 replicates (FDR=1−PPV). To combine both PPV and TPR, we use the F1 score (Van Rijsbergen 1979)

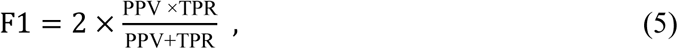

which is the harmonic mean of PPV and TPR, to measure the correct detection rate. A high F1 score implies that PPV and TPR are both high and balanced. The average detected hotspot number, F1 score and FDR are together used to assess the performance of methods, and a quality method should have the ability to provide the result with correct hotspot numbers, high F1 score and low FDR in hotspot detection.

Figure 5 depicts the F1 scores (y-axis), FDRs (x-axis) and average detected hotspot numbers (the numbers in the brackets) with the different thresholds over the 1000 simulation replicates. The average threshold of the Q-method is 7.58 (*β*=7.58). Using this threshold, the average number of detected hotspot is 9.94 (the true number is 6), indicating greater possibility of detecting extra false hotspots, and the associated F1 score and FDR are 0.749 and 0.400, respectively. With the proposed statistical procedure using the EQF-bin permutation, the average values of the thresholds *γ*_1,0.05_, *γ*_2,0.05_, …, *γ*_7,0.05_ and *γ*_8,0.05_ for qFreq(1), qFreq(2),…, qFreq(7) and qFreq(8) are 25.63, 18.45, 14.05, 11.53, 10.07, 9.11, 8.40 and 7.84 (≅ *β*), respectively, and the average numbers of hotspots detected under these thresholds are 1.59, 3.11, 4.46, 5.60, 6.57, 7.53, 8.52 and 9.46, respectively. The average hotspot number detected by using *γ*_4,0.05_ is 5.60, which is closest to the true number 6. The associated F1 scores are 0.419, 0.681, 0.836, 0.896, 0.891, 0.859, 0.814 and 0.770, respectively, and the associated FDRs are 0.000, 0.001, 0.020, 0.071, 0.148, 0.228, 0.306 and 0.371, respectively. In Figure 5, the closer a result with a threshold is to the upper left corner, the better it performs, simply because the F1 scores are higher and the FDRs are lower. Apparently, as compared to the result of the Q-method with the threshold value *β*=7.58, the results with *γ*_3,0.05_, *γ*_4,0.05_, *γ*_5,0.05_, *γ*_6,0.05_, *γ*_7,0.05_ and *γ*_8,0.05_ are closer to the upper left corner, and those with *γ*_1,0.05_ and *γ*_2,0.05_ are farther from the upper left corner. The results with *γ*_4,0.05_ and *γ*_5,0.05_ are better as they provide closer average hotspot numbers (5.60 and 6.57) to the true number 6, lower FDRs (0.071 and 0.148) and higher F1 scores (0.896 and 0.891), and the best result is obtained with *γ*_4,0.05_, not with *γ*_6,0.05_, in this specific setting (see CONCLUSION AND DISCUSSION for the reason). As expected, the Q-method produces a liberal threshold, which serves as the lower bound for the series thresholds *γ*_*n*,0.05_’s, and detects more spurious hotspots. The proposed procedure can provide much stricter thresholds, some of which may yield better results with correct hotspot numbers, higher F1 scores and less spurious hotspots (lower FDRs), for the assessment. The same results are also obtained with the QTL-interval permutation (not shown).

**Figure 5.**
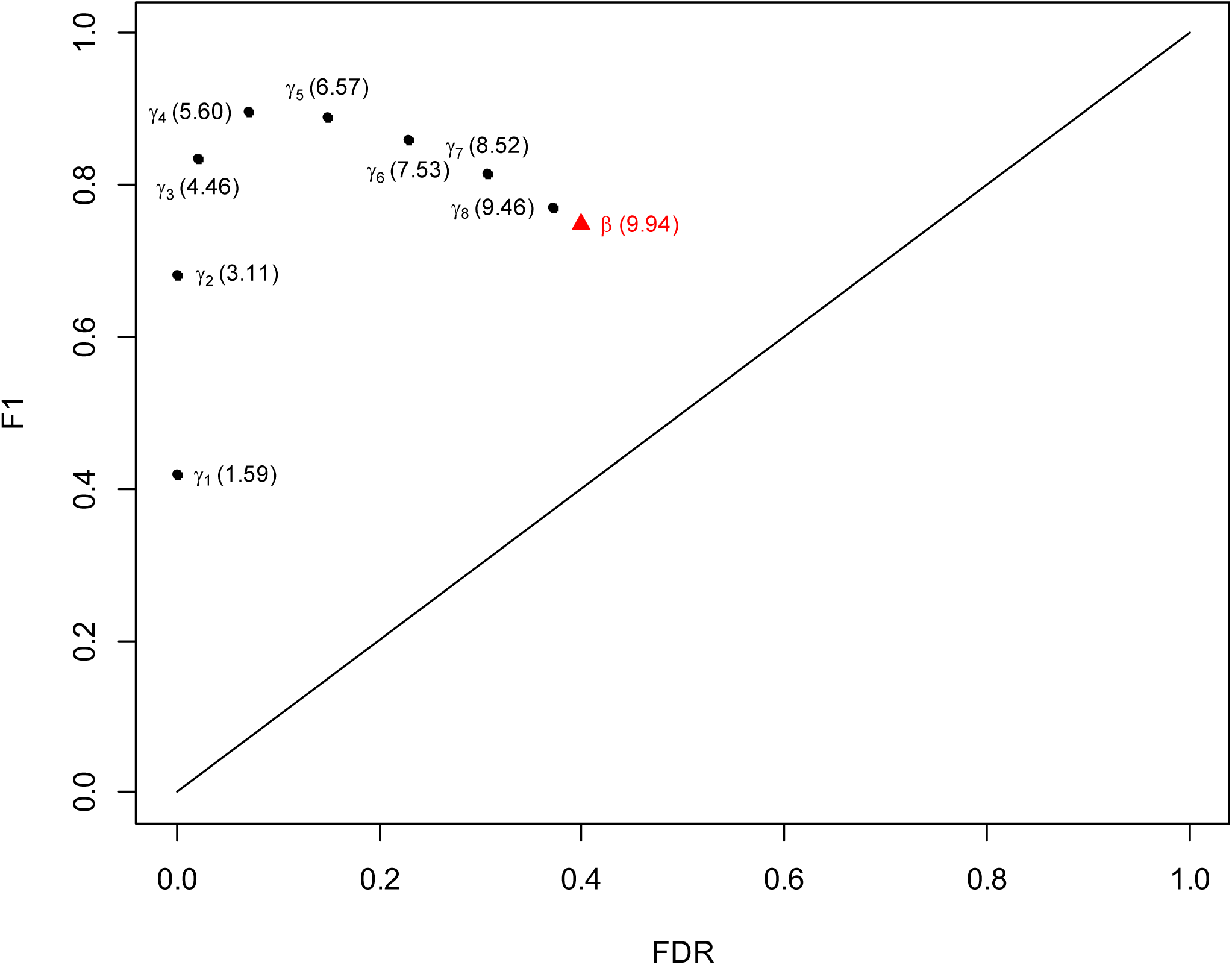
The F1 scores (y-axis), FDRs (x-axis) and average detected hotspot numbers (the numbers in the brackets) of the hotspot detection with the different thresholds over the 1000 simulation replicates. The dots denote the average thresholds *γ*_n,0.05_, *n* = 1,2, …, 8, obtained by the proposed statistical procedure and are 25.63, 18.45, 14.05, 11.53, 10.07, 9.11, 8.40 and 7.84, respectively. The triangle denotes the average threshold *β* obtained by the Q-method and is 7.58. The true hotspot number is 6.

#### Genetical genomics experiments

For the third part, concerning the information loss, we mimic the simulation study in Nato *et al.* (2012) with modification to simulate a small-scale genetical genomics data set, and then apply the proposed statistical procedure, Q-method, Breitling’s method (the N-method) and Neto’s method (the NL-method) to analyze the data set for evaluation. The data set contains 100 backcross progeny with 5 chromosomes of length 100 cM and 600 molecular traits. Each chromosome contains 50 equally spaced markers. The 600 traits are assumed to belong to three different trait categories. Three hotspots are considered: (1) A small hotspot A is cause by a gene at 50 cM on the 1^st^ chromosome. The gene controls 100 traits with heritabilities 0.3~0.4 showing moderate to high LOD scores (see Figure S3A) in QTL mapping. The 100 pleiotropic traits are assigned to the first trait category; (2) A big hotspot B is caused by a gene located at 50 cM on the 3^rd^ chromosome. The gene influences 300 traits belonging to the second category. Among the 300 pleiotropic traits, half have heritabilities 0.1~ 0.45 showing small to high LOD scores (see Figure S3B), and half have heritaibilities 0.3~ 0.45 showing moderate to high LOD scores (see Figure S3B); (3) A big hotspot C is caused by a gene at 50 cM on the 5^th^ chromosome. The gene controls 200 traits that belong to the third category. The heritabilities of the 200 pleiotropic traits are 0.1~0.2 showing small LOD scores (see Figure S3C). The pairwise correlations between traits are shown in Figure S3D. The bin size of 2 cM (Δ=2 cM) is used in the analysis (similar to that in Neto et al.). The bin containing the estimated QTL position will be given to 1 (and 0 otherwise) to construct the QTL matrix. Both the proposed statistical procedure and the Q-method operate on the QTL matrix for detection analysis. The trait grouping in the proposed procedure considers that all pleiotropic traits are correctly allocated to the same categories (perfect trait grouping). For the N-method and NL-method, we follow Neto *et al.* (2012) to adopt 1.5-LOD support intervals for the backcross to decrease the spread of the hotspots.

Figure 6 shows the results of the four methods for the simulated data set. Figure 6A presents the hotspot architecture constructed using a single-trait LOD threshold of 2.47 and the 5% significance hotspot size thresholds obtained by the Q-method, N-method, and the proposed procedure. The permutation thresholds delivered by the Q-method and N-method are 3 and 11, respectively. Under the thresholds, there are 18 and 38 significant bins (detected hotspots), respectively (see Figure 6A), showing that, in addition to detecting the three true hotspots, several spurious hotspots are also detected near the true hotspots. The permutation thresholds obtained by the proposed procedure for assessing the significance of at least 1, 2, 3 and 4 spurious hotspots are *γ*_1,0.05_ =208, *γ*_2,0.05_ = 58, *γ*_3,0.05_ = 47 and *γ*_4,0.05_ = 34, respectively. Under these thresholds, there are 1, 1, 4 and 5 detected hotspots, respectively, indicating that less spurious hotspots are detected due to higher thresholds. Using *γ*_3,0.05_ = 47 as a threshold, the three true hotspots (the highest bins on chromosomes 1, 3, and 5) and one spurious hotspot (right next to the true hotspot on chromosome 3) are significant. Figures 6, B-F, present the hotspot architectures inferred using the NL-method LOD thresholds of 5.36, 3.04, 2.03, 1.24, and 1.07 that aim to control GWER of 5% for spurious hotspots of sizes 1, 5, 20, 77, and 100, respectively. Under these thresholds, there are 31, 22, 11, 3 and 1 significant bins (hotspots). Using LOD thresholds of 5.36, 3.04, 2.03, not only the three true hotspots but also some spurious hotspots (the secondary peaks around the true hotspots) are detected. Using a LOD threshold of 1.24, only the three true hotspots (on chromosomes 1, 3, and 5) are significant. Using a LOD threshold of 1.07, only the true hotspot on chromosome 3 is detected as significant. The above shows that, if trait grouping is perfect, the proposed statistical procedure is applicable and can obtain comparable results in QTL hotspot detection in the genetical genomics experiments. But note that if trait grouping is not perfect, the hotspot thresholds decrease and the possibility of detecting more spurious hotspots increases (not shown).

**Figure 6.**
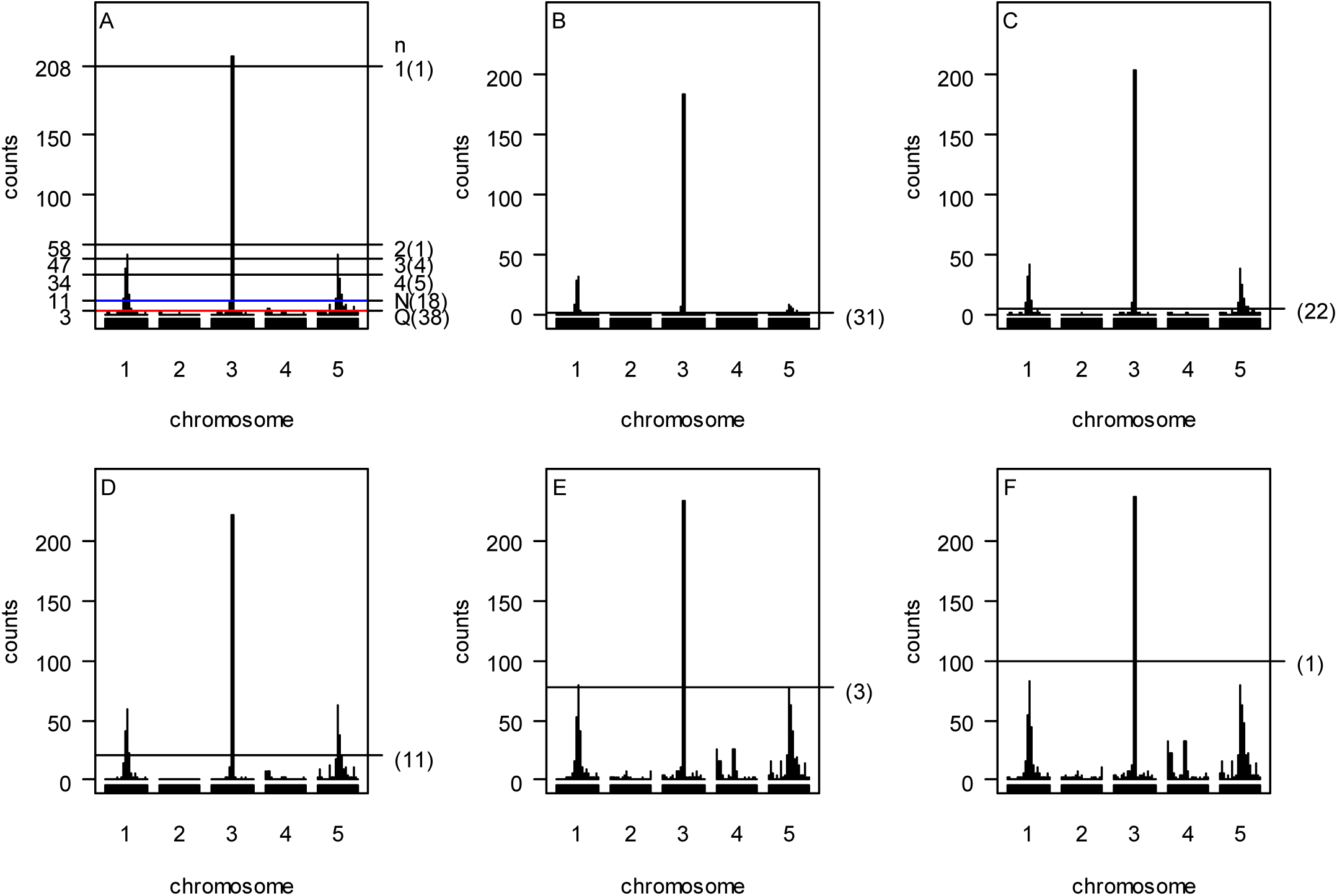
Panels (A-F) The proposed statistical procedure, Q-method, N-method and NL-method analyses for simulated example. Panels (A) Inferred hotspot architecture using a single-trait permutation LOD threshold of 2.47 corresponding to a GWER of 5% of falsely detecting at least one QTL somewhere in the genome. The hotspots on chromosomes 1, 3 and 5 have sizes 50, 210, and 50, respectively. The blue line at count 11 corresponds to the hotspot size threshold at a GWER of 5% according to the N-method. The red line at count 3 corresponds to the Q-method’s 5% significance threshold. The thresholds y_l,0.05_, y_2,0.05_, y_3,0.05_ and y_4,0.05_ obtained by the proposed procedure are 208, 58, 47 and 34, respectively. Panels (B, C, D, E and F) Hotspot architecture inferred using different permutation thresholds by the NL-method; Hotspot architectures computed using QTL mapping LOD thresholds of 5.36 (B), 3.04 (C), 2.03 (D), 1.24 (E), and 1.07 (F) that aim to control GWER at a 5% error rate for spurious QTL hotspots of sizes 1, 5, 20, 77, and 100, respectively. The number in the bracket is the number of detected hotspots. Results are based on 1000 permutations. Q: The Q-method; N: The N-method.

To sum up, the simulation study shows that the proposed statistical procedure can control GWERs at the target levels for the QTL data with correlation structure, has the ability to produce quality results by offering a sliding scale of thresholds from high to low for QTL hotspot detection, and is applicable to the hotspot analysis in genetical genomics studies.

## CONCLUSION AND DISCUSSION

Both genetical genomics experiments and public QTL databases can provide abundant QTLs for genome-wide detection of QTL hotspots to explore the genetic mechanism of quantitative traits in biological studies. A single genetical genomics experiment can produce an adequate individual-level data set that contains the genotypes and a large enough number of phenotypes for QTL mapping and further for QTL hotspot detection. On the contrary, public QTL databases consistently collect summarized QTL data for many phenotypic traits from numerous independent experiments that allows for detection of QTL hotspots. Several methods, mainly for using individual-level data, have been proposed to detect QTL hotspots (see introduction). In this article, we develop a statistical procedure for detecting QTL hotspots at the genome-wide level by using summarized QTL data in public databases. We first obtain the QTL intervals from public databases and use them to compute the EQF matrix for operation. We then derive a permutation algorithm on the EQF elements or the QTL intervals to compute a sliding scale of EQF thresholds that range from conservative to liberal for assessing the significance of QTL hotspots. To consider the correlation structure among traits, the correlated traits are grouped together as a trunk or unit of permutation to obtain stricter thresholds. As shown in simulation study, with the grouping strategy, our statistical procedure can control the GWERs of the test statistic at the target levels under varying strengths of correlation among QTLs and can provide much more rigorous thresholds for hotspot detection with higher correct (lower incorrect) detection rates. Besides, to evaluate the information loss between the two types of data in hotspot detection, we apply the proposed procedure, Q-method, Breitling’s method and Neto’s approach to analyze a simulated genetical genomics data set and compare their results. It shows that our procedure is comparable to the methods using individual-level data if the pleiotropic (correlated) traits can be all grouped together into the same trait categories. In the GRAMENE rice database, more than 100 QTL hotspots were detected in the genome. We also conduct a genome-wide comparative analysis of the detected hotspots and the 122 known genes in the Rice Q-TARO database. The comparative analysis shows that the QTL hotspots and genes are conformable in the sense that they co-localize closely and are functionally related to the associated traits. An R package of our proposed statistical procedure called QHOT is available on http://www.stat.sinica.edu.tw/~chkao/ and is being submitted to Comprehensive R Archive Network (CRAN). The R codes can readily produce both numerical and graphical outputs that would allow visualization of several features, including the EQF architecture, the QTL components of the hotspots, and nearby known genes, at the genome-wide level (see Figures 2, 3 and S2) for exploring the interplay among QTLs, genes and traits.

The QTL hotspot detection using public databases relies on the great number of QTLs collected from numerous independent QTL mapping studies. The collected QTLs are usually mapped for various traits and detected by different statistical tools under a wide range of experimental parameter settings. Quite often some biologically interesting and popular important traits are more frequently investigated and mapped for QTLs in the studies, resulting in a group of traits strongly mapping to the same or closely linked QTLs mostly due to the genetic correlations among them. Take rice as an example, the agronomic traits, such as panicle numbers, grains per panicle and grain weight, plant height, heading time, grain quality, insect and disease resistant, sterility, etc., are often investigated and highly correlated to each other (Swamy *et al.* 2011; Trijatmiko *et al.* 2014; Wu *et al.* 2016). To account for the correlation structure among traits, we group and permute these traits together to determine a series of thresholds for hotspot detection. Other grouping, such as by the already known correlations between traits or biological prior knowledge on traits, is also applicable. The simulation study shows that grouping of highly correlated traits is effective to control the error rates of falsely detecting a hotspot. If genetically correlated traits are not grouped together and used as control, the error rates may inflate greatly and will be higher than the target levels. The error rates are not sensitive to the situation in which uncorrelated traits are falsely grouped together. Ideally, we would like to have a perfect grouping, in which traits are correlated in the same categories and uncorrelated in different categories (the first and third parts of simulation), or the pleiotropic traits are all grouped into the same trait categories (the third part of simulation), to cope with their correlation features. However, in practice, a perfect grouping is not always possible because correlations among traits are likely to be common, and the pleiotropic traits may be allocated to different categories. For example, the yield and disease resistant are correlated and assigned into different categories in GRAMENE database. The grain weight and plant architecture are pleiotropic traits and classified into different categories (Fujita *et al.* 2013; Xu *et al.* 2015). The GRAMENE rice database classifies all traits into nine major trait categories based on the general agronomic consideration, which is imprecise in the sense that traits in the same categories may have different strengths of genetic correlation, and traits in different categories still preserve certain degrees of genetic correlation. This allows us to argue that underestimation of the threshold and the problem of information loss are still very likely to occur to some extent in the real example analysis, resulting in excesses of observed over expected hotspot numbers (as validated in the real example analysis and simulation study).

In the QTL hotspot analysis using genetical genomics experiments, the proposed statistical procedure can be extended as follows: First, QTL mapping is performed to obtain the QTL matrix; Second, the correlation coefficients among the traits are computed for grouping reference; Next, the QTLs for the (highly) correlated traits are grouped and permuted together to obtain a series of thresholds for assessing hotspot significance. Note that the trait grouping can be also done in different ways, such as by principal component analysis or cluster analysis (Abdi and Williams 2010; Everitt *et al.* 2011). Such a procedure is easy to implement and very cheap in computation as compared to the methods by permuting the original individual-level data in the hotspot analysis (see ***The permutation algorithm)***. It has been observed that QTLs or genes for genetically correlated traits have a tendency to cluster on the same or adjacent regions of chromosomes in several organisms, which may be due to linkage, pleiotropy or natural selection for co-adapted traits (Studer and Doebley 2011; Wu *et al.* 2015). The grouping strategy attempts to take the clustering phenomenon into account in the detection of QTL hotspots. In data collection, it would be important to collect as many QTLs and genes for various traits as possible of to identify the clustering phenomenon and explore the interplays among QTLs, genes and traits. Our real example analysis considers the 8216 QTLs for 236 component traits in GRAMENE rice database for hotspot detection and compares with the 122 known genes in QTARO database. The results display the clustering phenomenon of QTLs and genes around the detected hotspots. For example, the well-known pleiotropic gene *SCM2* responsible for lodging resistance and yield is very close to the QTL hotspot [6,115.5-116] (see Figure S2), which contains QTLs for yield, vigor and development. Also, five closely linked genes, *Rc*, *qSD7-1/qPC7*, *OsHMA3*, *qCDT7* and *Ghd7*, with similar functions are localized around the detected hotspots [7,44.5-45] and [7,50-50.5]. Understanding the genetic architectures of quantitative traits at the genome-wide level has been a key and challenging issue in various areas of genetics, gene and genomics studies. This would rely heavily on effectively integrating and analyzing the information on the QTLs and genes from the published literature. Currently, several well-known databases of important organisms (see INTRODUCTION section) have consistently collected the information and allow researchers access to their well-curated datasets, and, therefore, increasing numbers of QTLs and genes for a variety of traits are available and ready for further application. There is a need to develop statistical methods for mining the useful information and knowledge from these databases. By using public databases, we develop a statistical procedure for detecting QTL hotspots, perform comparative analysis with the known genes for validation, and also identify several potential QTL regions of genes (not shown). The R codes can present the EQF architectures for hotspots (QTLs), genes and quantitative traits region by region along the genome to overview their connections. Our approach can provide a way to explore networks among QTL hotspots, genes and traits for dissecting the genetic architecture of complex traits in broad areas of biological studies.

---

**Figure S1.**
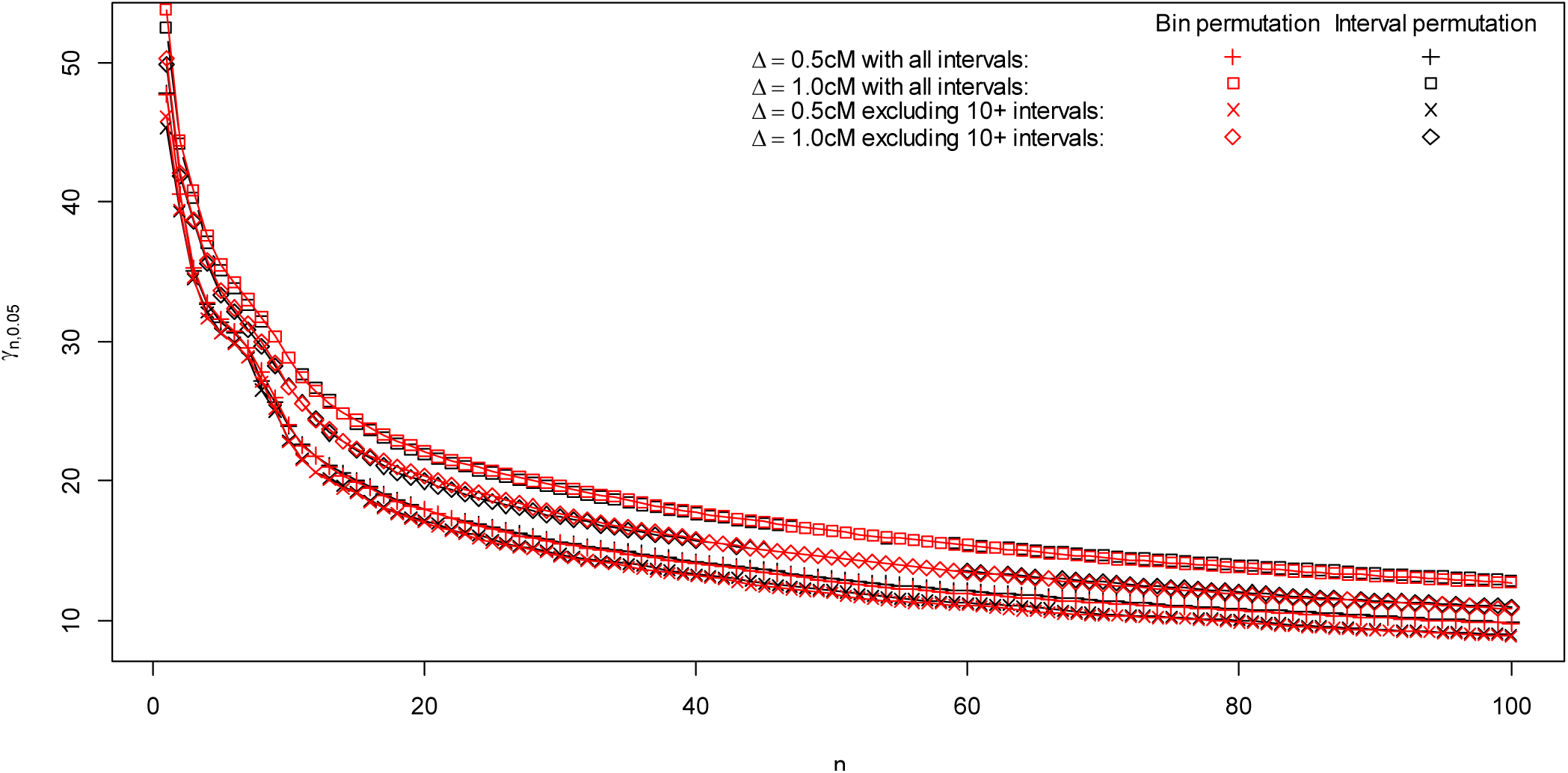
In order to explore the effects of bin sizes and QTL interval sizes, the thresholds of *γ*_n,0.05_, n=1, 2,…, 20, for using two kinds of bin sizes, 0.5 and 1 cM, with all intervals or removing the >10 cM, respectively are analyzed. In general, as bin sizes increase, the hotspot resolution will decrease, and the EQFs and threshold values will increase. Besides, these 10+ cM QTL intervals effect hardly to the detection results, since they contribute little to the EQF value. Finally, the thresholds obtained by permuting the EQF bins and the QTL intervals are very similar

**Figure S2.**
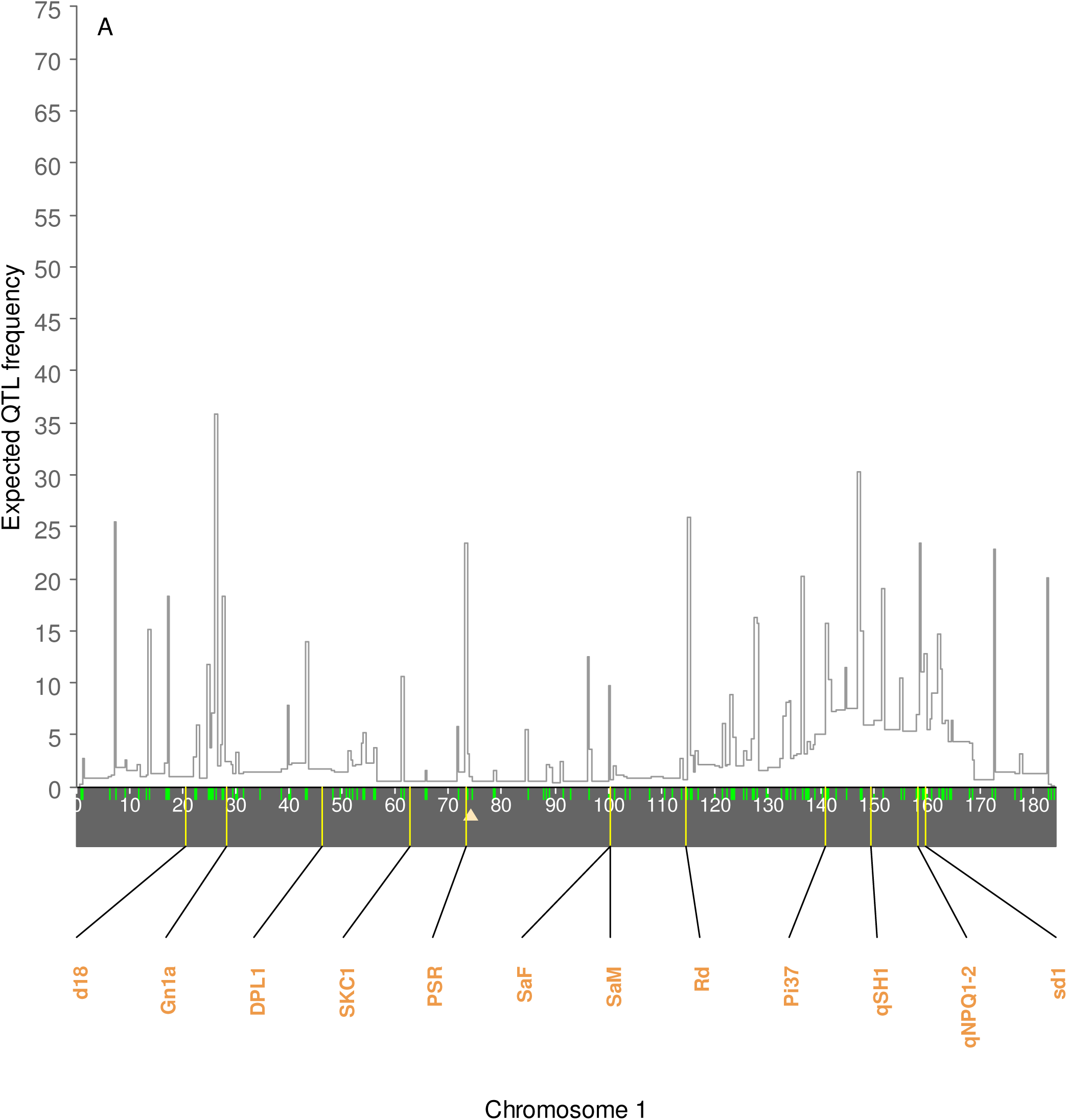
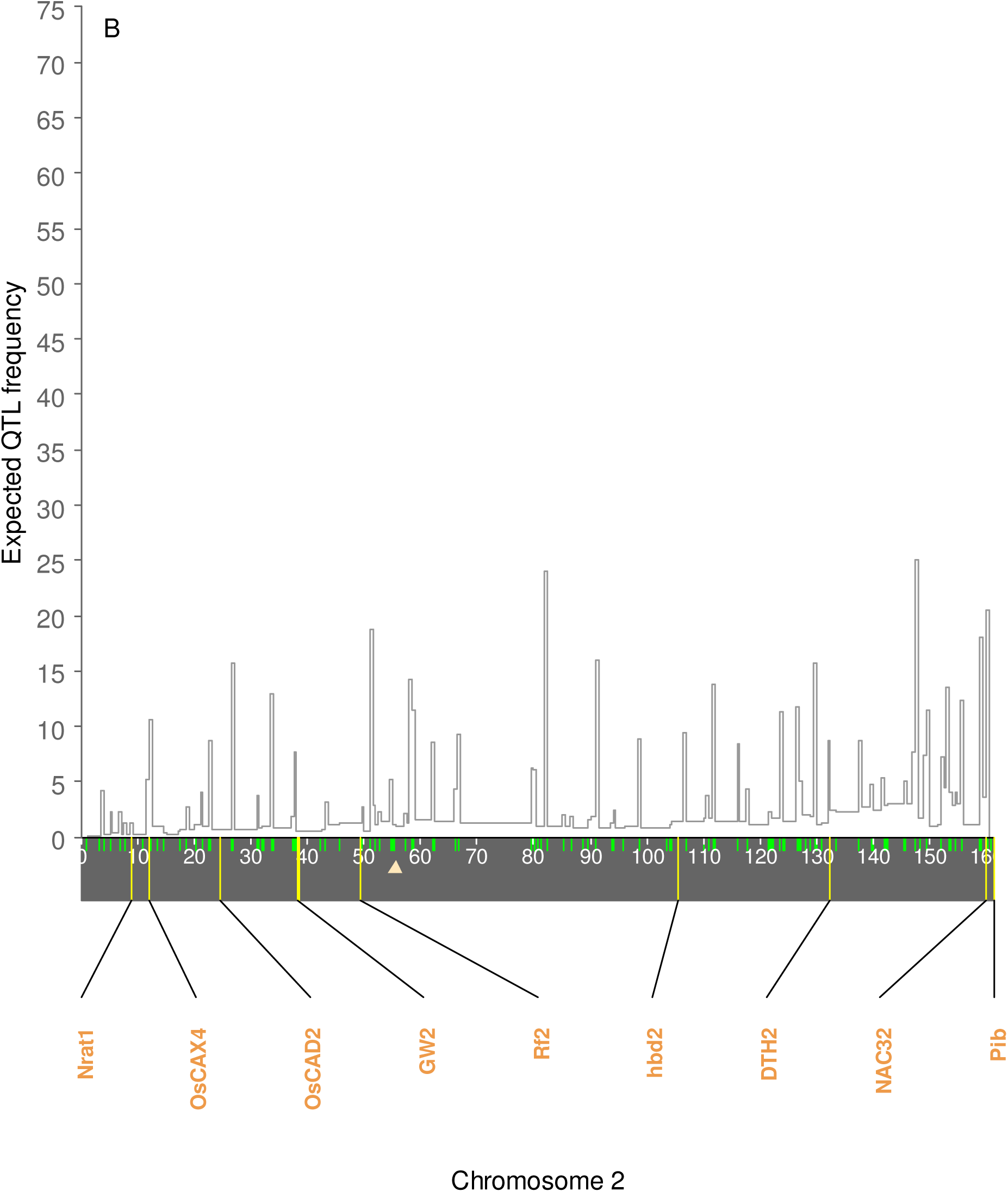
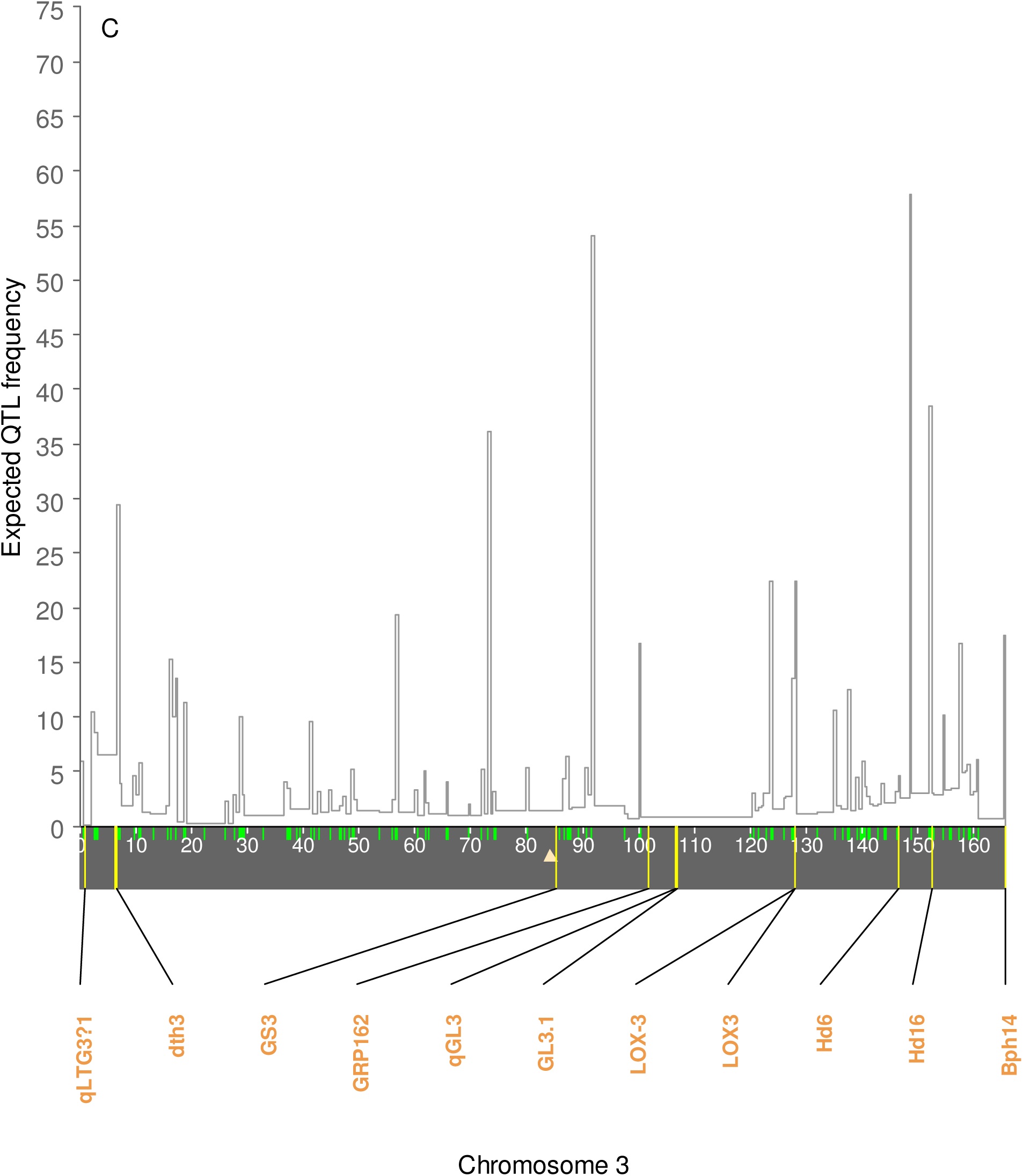
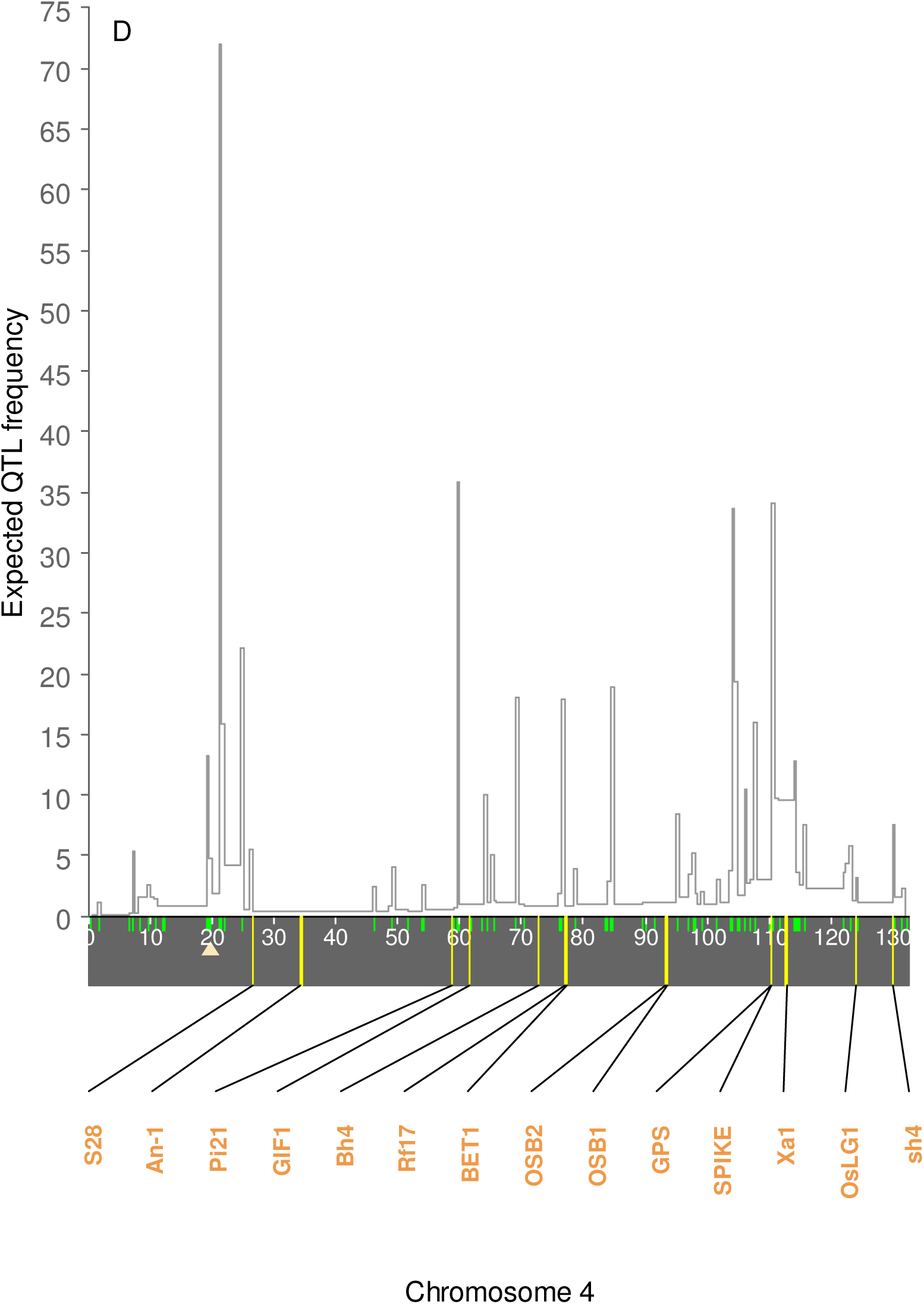
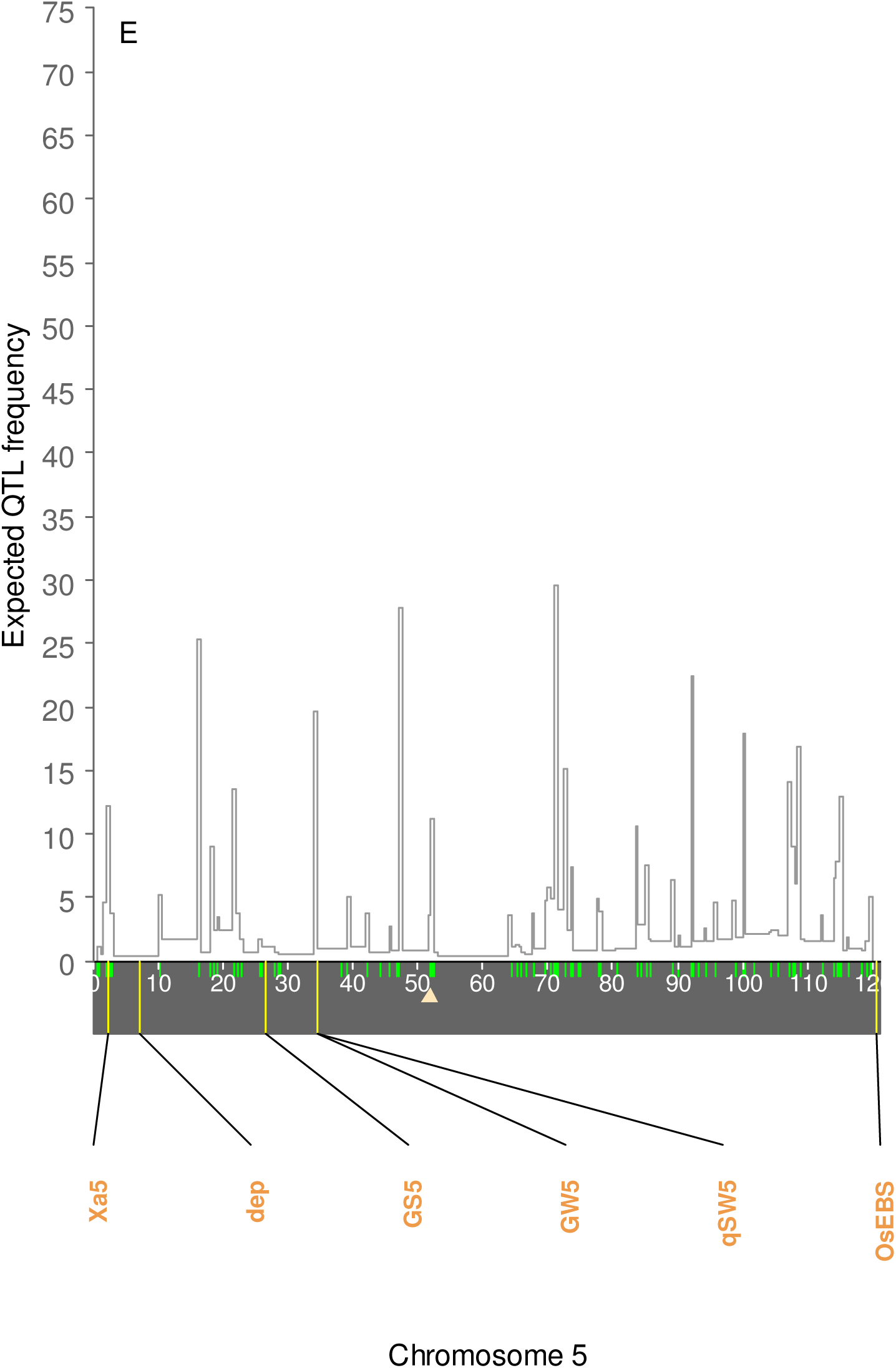
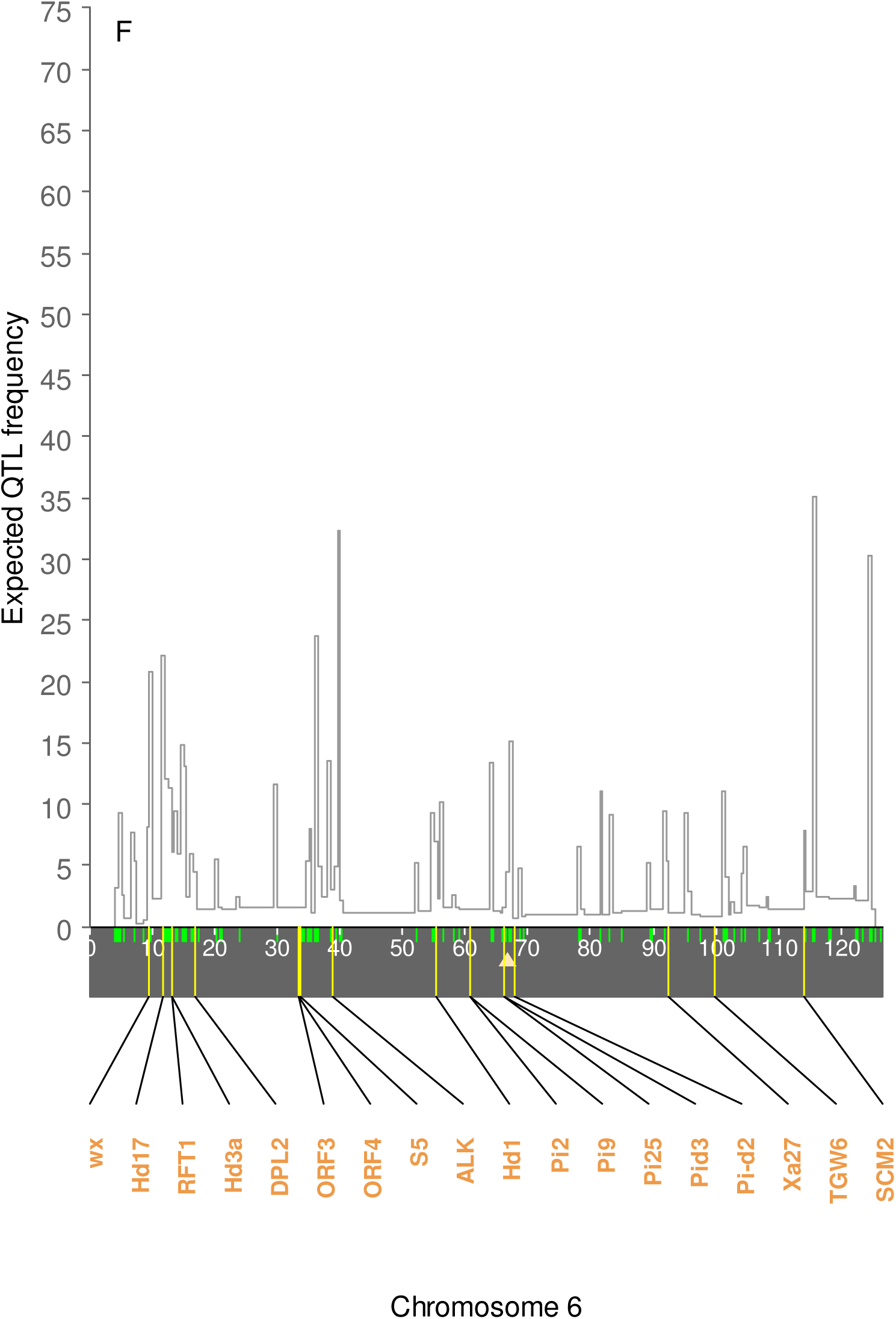
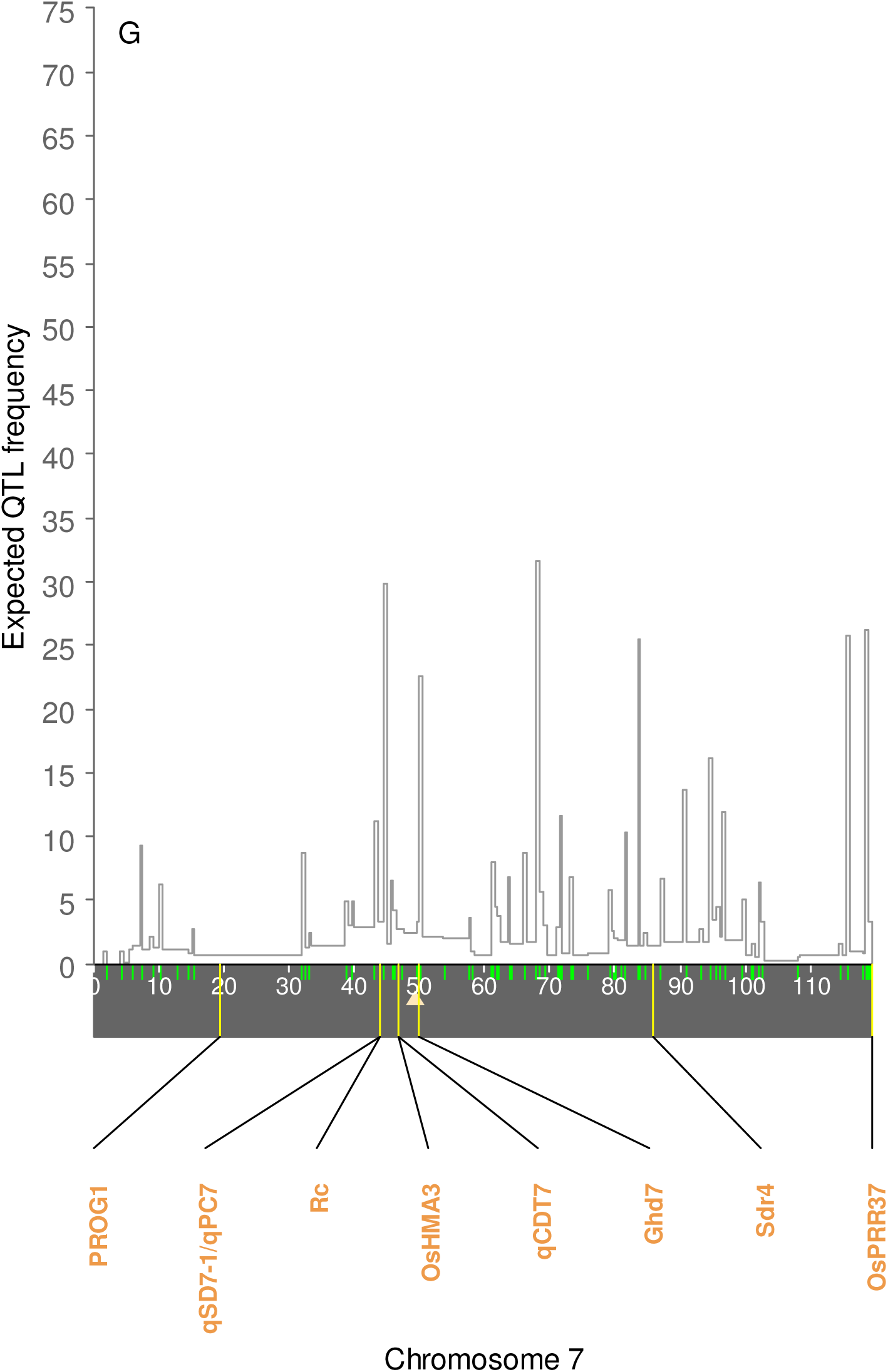
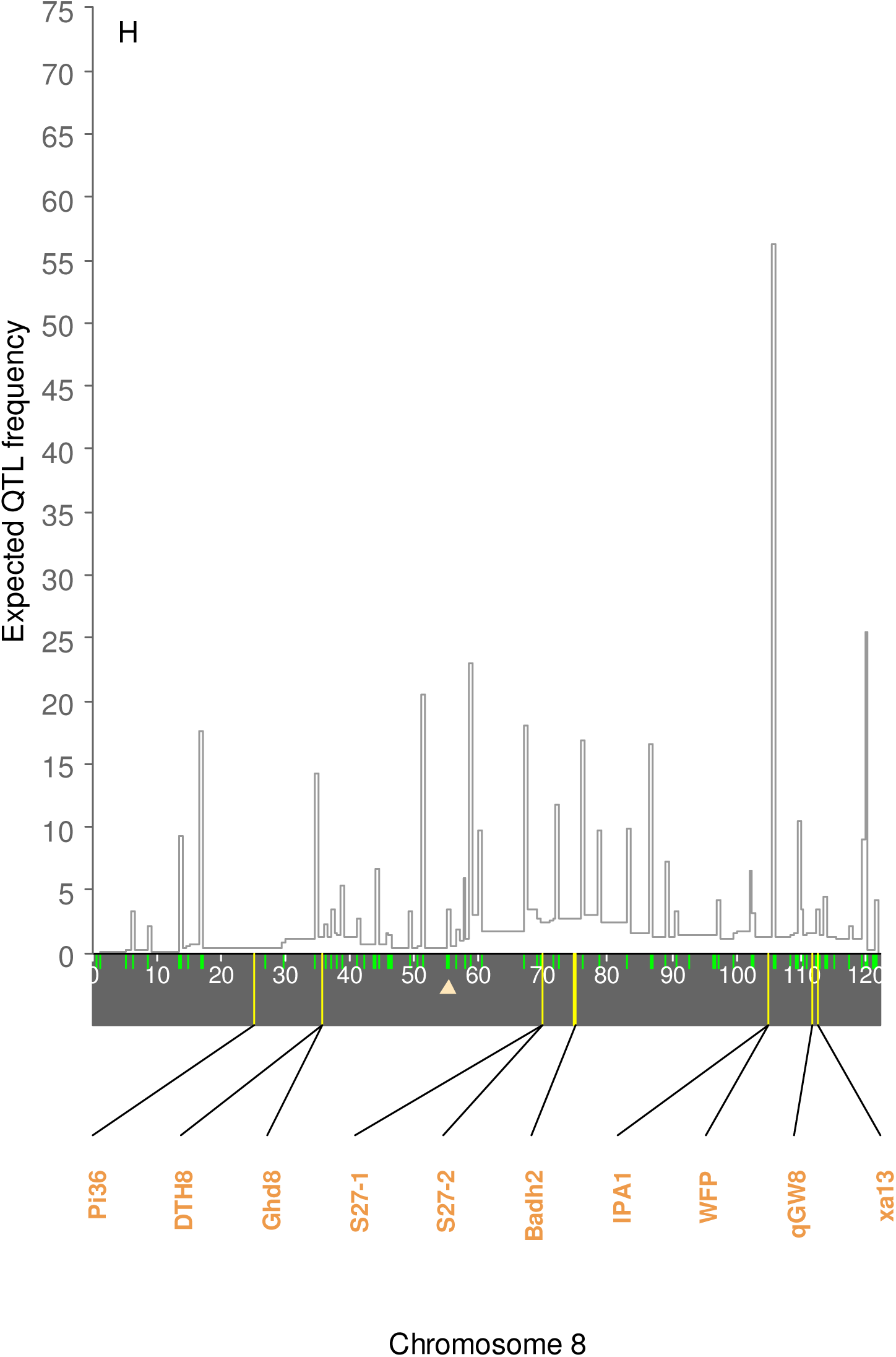
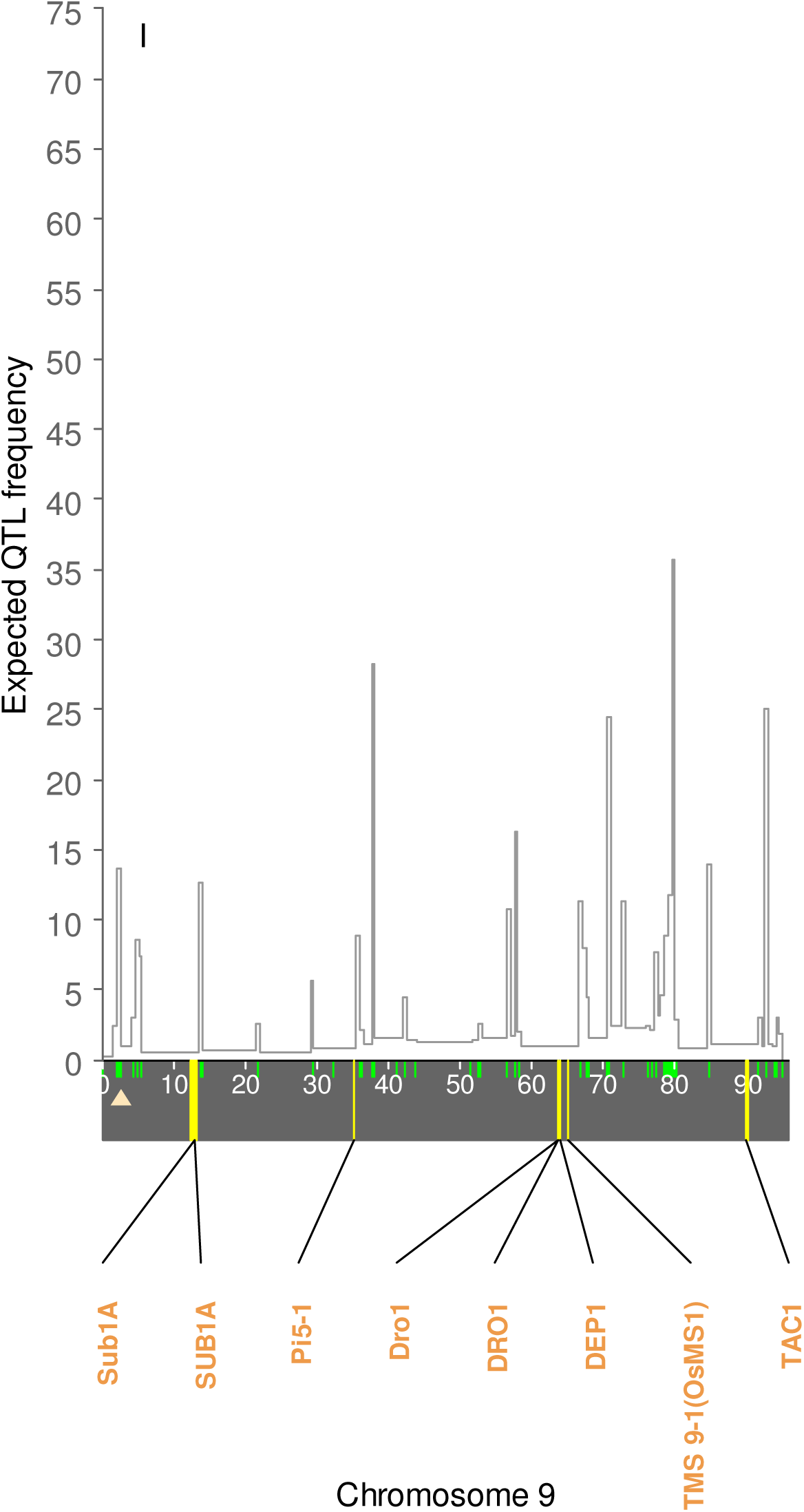
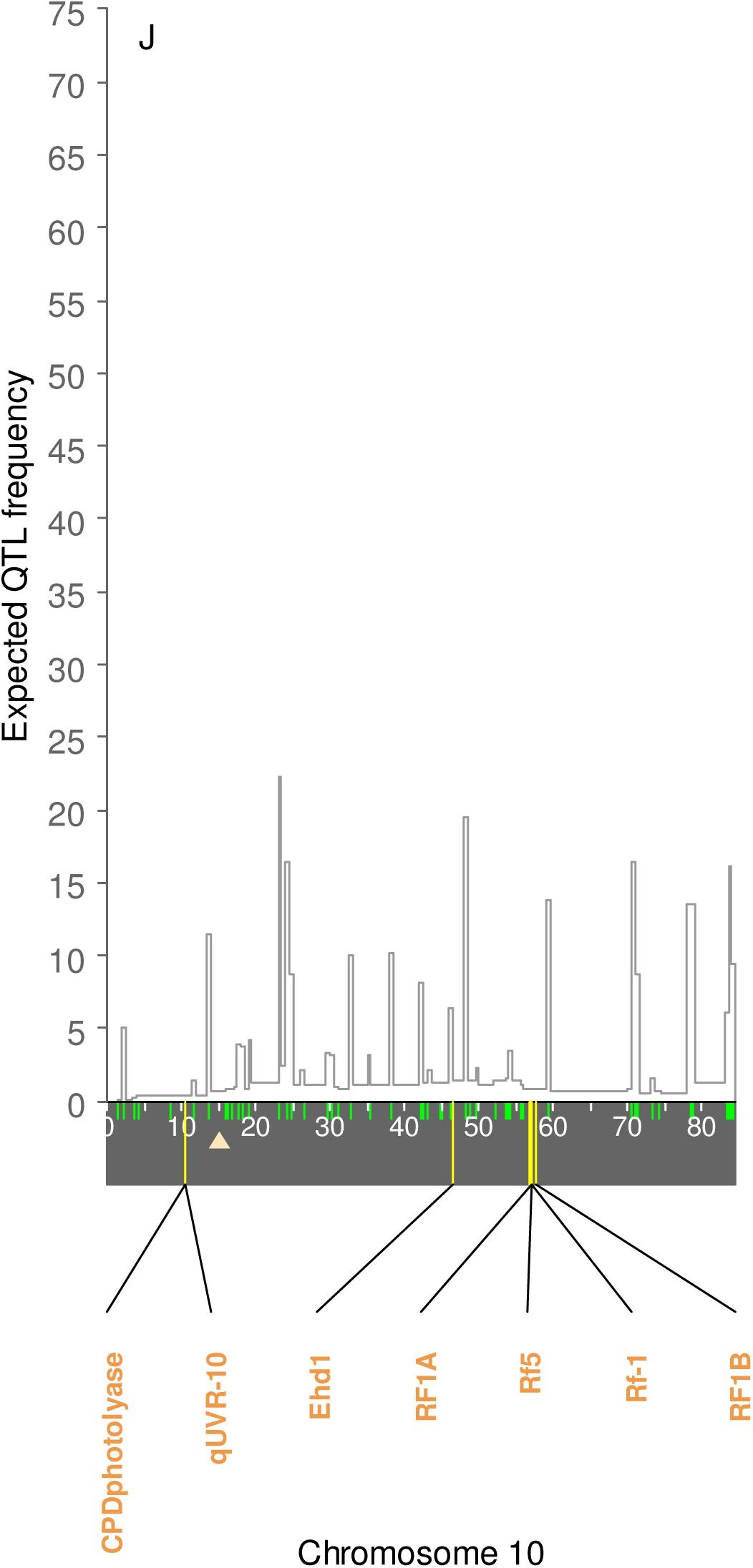
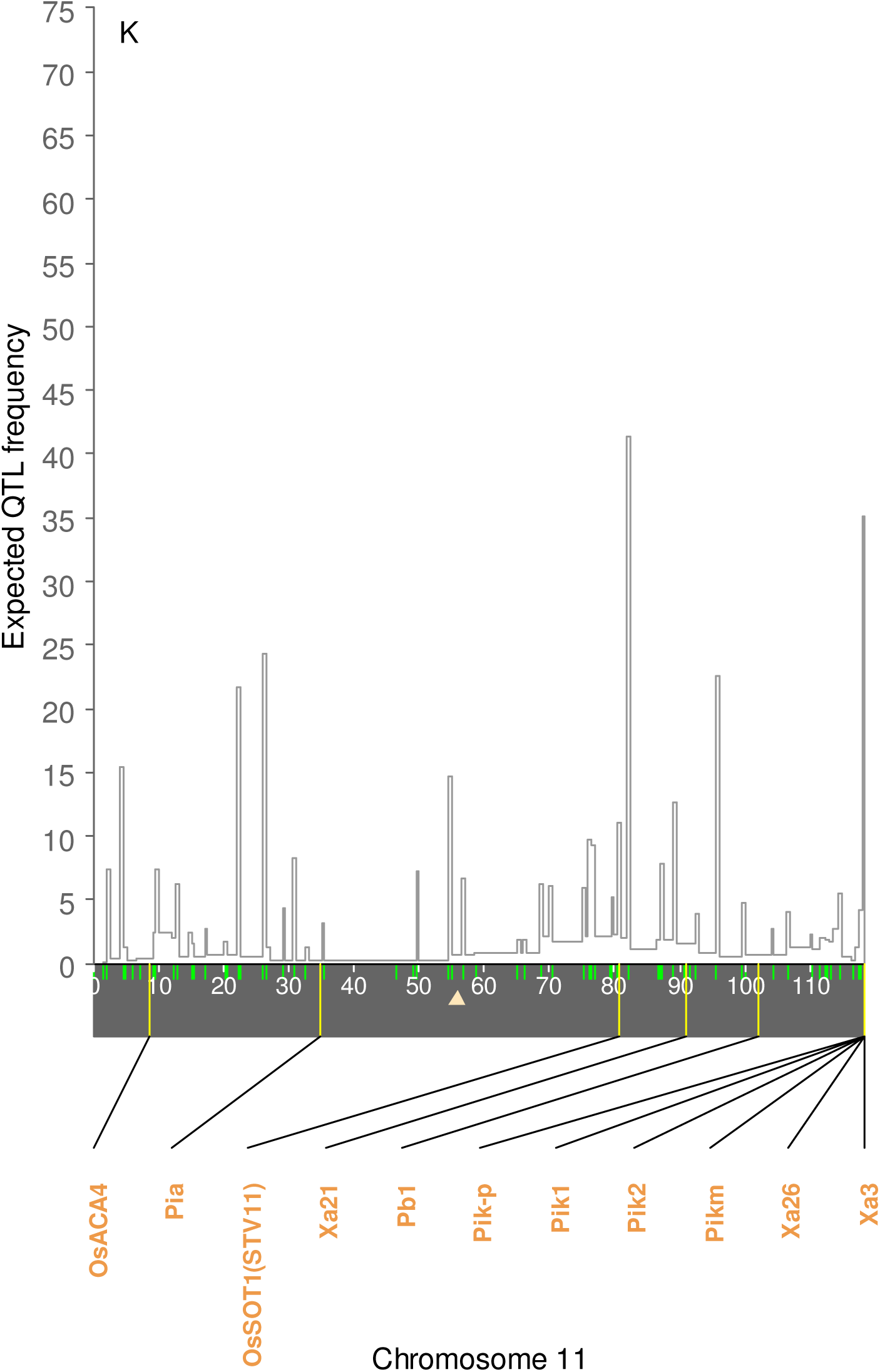
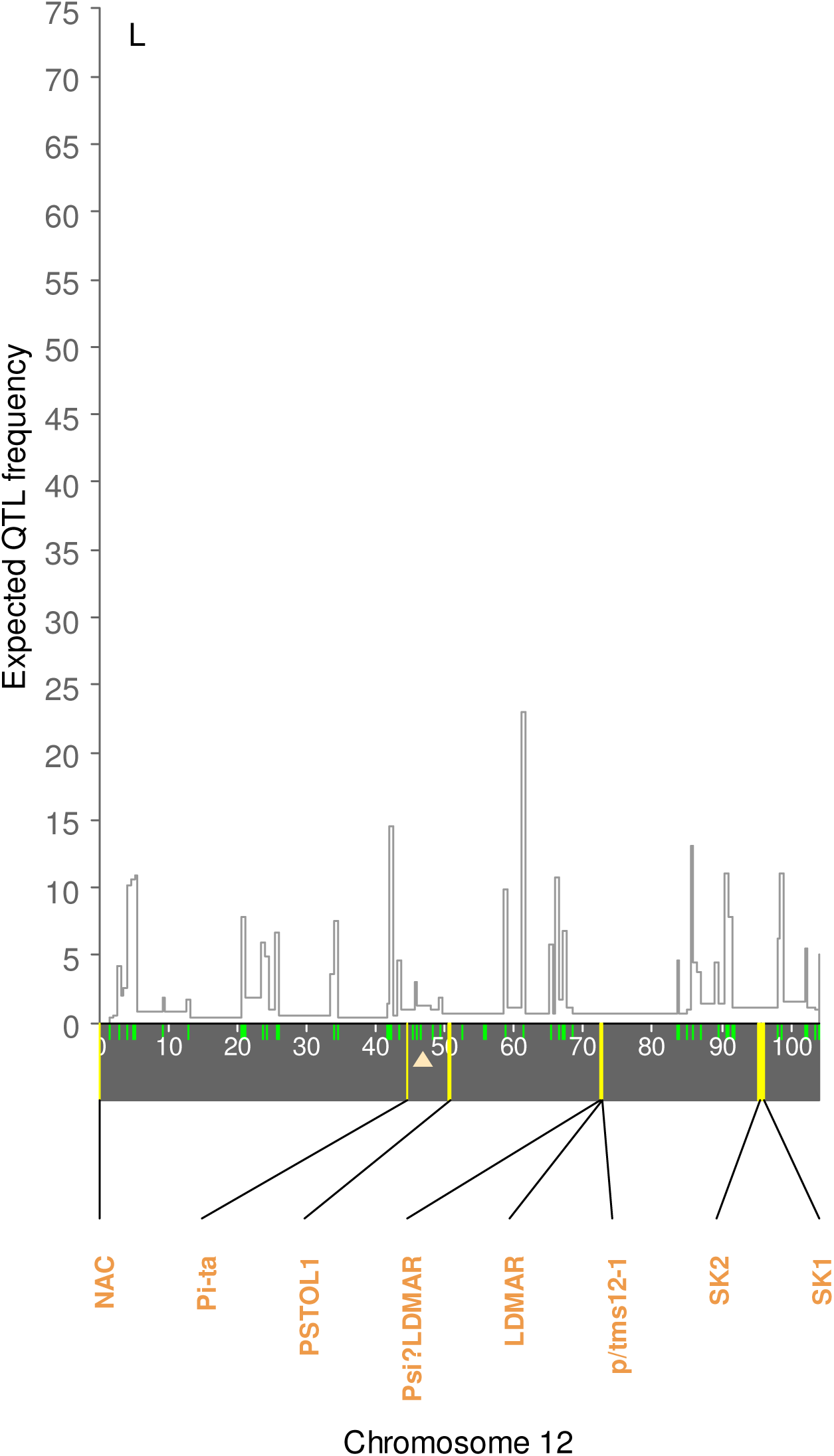
The EQF architectures and known gene 40 locations in the rice 12 chromosomes (A-L). The EQF architectures are constructed by using Gramene Rice database (http://www.gramene.org/), and the known genes (identified by natural variation analysis) are collected in Q-TARO database (http://qtaro.abr.affrc.go.jp/). The y-axes are the expected QTL frequencies. The green and yellow ticks on the x-axes denote the positions of the 1914 markers and 122 known genes.

**Figure S3.**
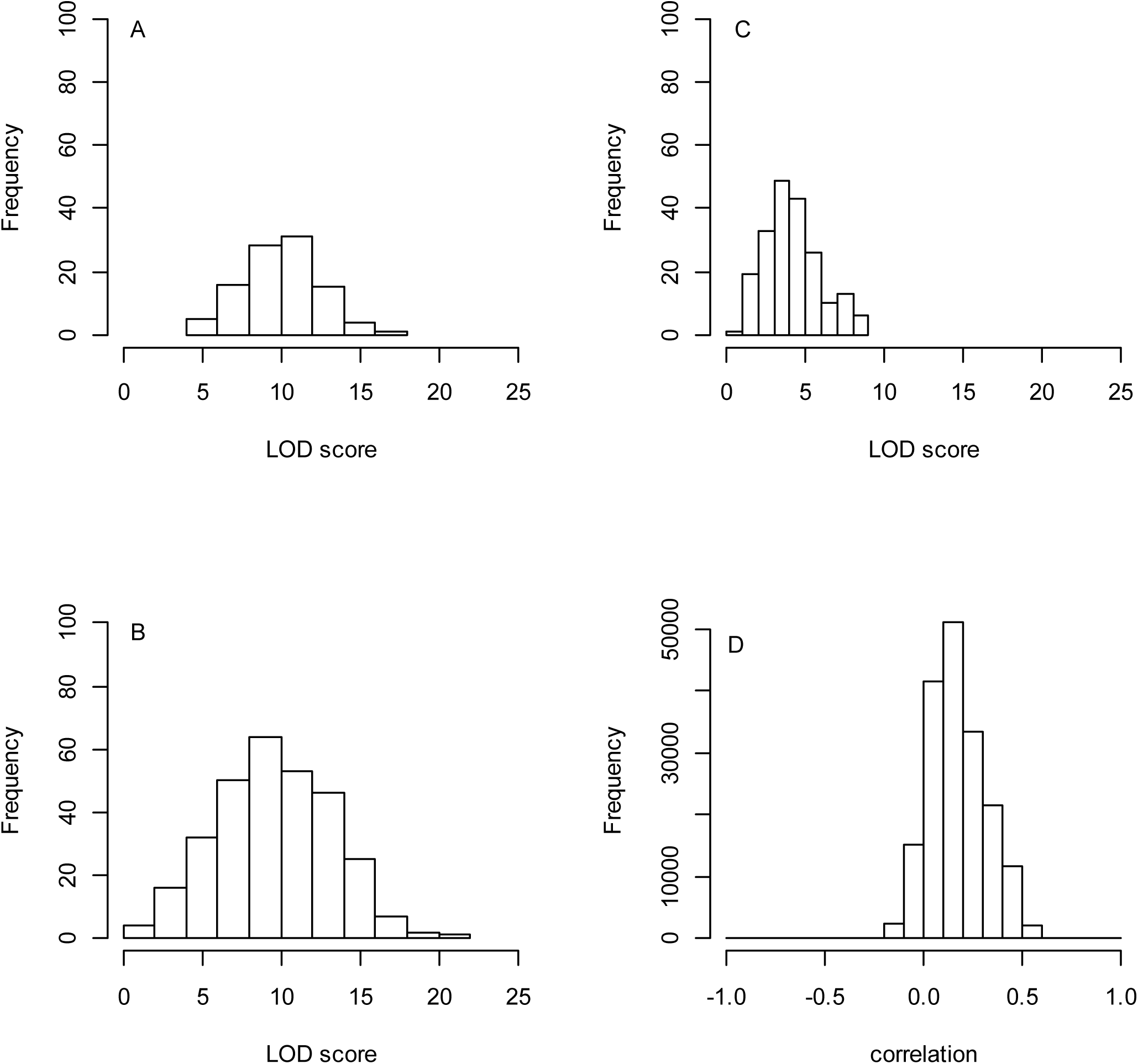
Distributions of hotspot LOD scores and pairwise correlations among traits for simulated genetical genomics data. Panels (A), (B) and (C) show the LOD score distribution for the hotspot on chromosome 1, chromosome 3, and chromosome 5, respectively. The histograms show the distribution of the LOD scores of the traits composing the hotspot at the hotspot peak location. Panel (D) shows the distribution of the pairwise correlations among traits for the simulated data.

**Table S1.**
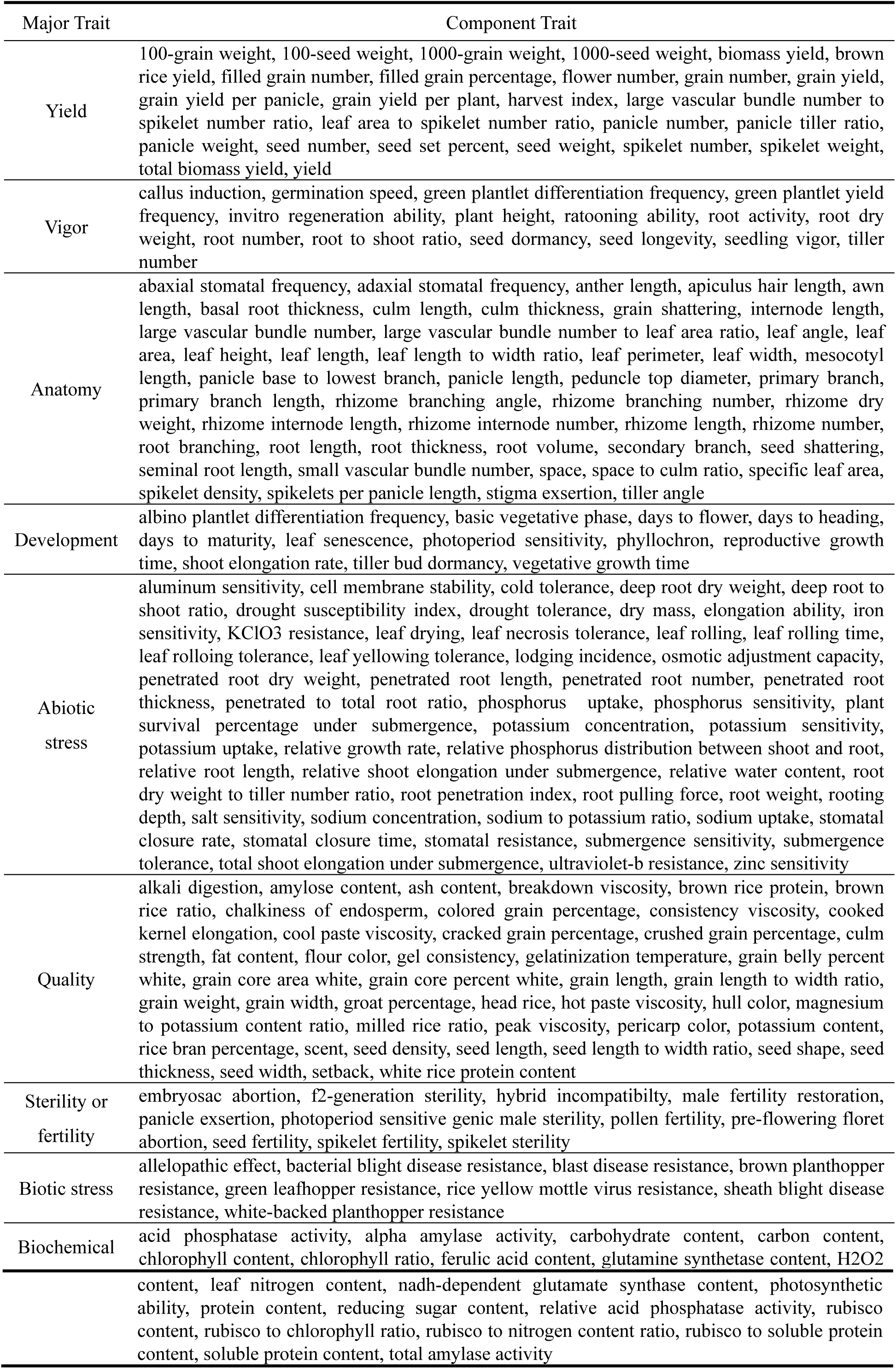
Nine major traits and their component traits in Gramene Rice database (http://www.gramene.org/).

**Table S2.** The 8216 QTLs responsible for the 9 major traits (see Table S1) in the 12 chromosomes from Gramene Rice database (http://www.gramene.org/).

| Chr. | Length(cM) | Yield |  | Vigor |  | Anatomy |  | Development |  | Abiotic stress |  | Quality |  | Sterility or fertility |  | Biotic stress |  | Biochemical |  | Total |  |
| --- | --- | --- | --- | --- | --- | --- | --- | --- | --- | --- | --- | --- | --- | --- | --- | --- | --- | --- | --- | --- | --- |
| 1 | 184.19 | 351<sup>a</sup> | <u>1.91<sup>b</sup></u> | 310 | <u>1.68</u> | 165 | <u>0.90</u> | 86 | 0.47 | 118 | <u>0.64</u> | 71 | <u>0.39</u> | 61 | <u>0.33</u> | 61 | <u>0.33</u> | 24 | <u>0.13</u> | **1247** | **<u>6.77</u>** |
| 2 | 161.49 | 198 | 1.23 | 159 | 0.98 | 128 | 0.79 | 95 | <u>0.59</u> | 92 | <u>0.57</u> | 44 | 0.27 | 28 | 0.17 | 44 | <u>0.27</u> | 19 | <u>0.12</u> | **807** | **5.00** |
| 3 | 165.83 | 206 | 1.24 | 257 | <u>1.55</u> | 149 | <u>0.90</u> | 152 | <u>0.93</u> | 69 | 0.42 | 73 | <u>0.45</u> | 75 | <u>0.45</u> | 33 | 0.20 | 11 | 0.07 | **1025** | **<u>6.18</u>** |
| 4 | 132.72 | 178 | <u>1.34</u> | 188 | <u>1.42</u> | 144 | <u>1.08</u> | 58 | 0.44 | 73 | <u>0.55</u> | 26 | 0.20 | 29 | 0.22 | 32 | 0.24 | 15 | <u>0.11</u> | **743** | **<u>5.60</u>** |
| 5 | 121.24 | 166 | <u>1.37</u> | 141 | <u>1.16</u> | 87 | 0.72 | 51 | 0.43 | 57 | 0.47 | 70 | <u>0.59</u> | 26 | 0.21 | 12 | 0.10 | 14 | <u>0.12</u> | **624** | **5.15** |
| 6 | 126.56 | 213 | <u>1.68</u> | 123 | 0.97 | 138 | <u>1.09</u> | 75 | <u>0.61</u> | 43 | 0.34 | 88 | <u>0.71</u> | 43 | <u>0.34</u> | 44 | <u>0.36</u> | 10 | 0.08 | **777** | **<u>6.14</u>** |
| 7 | 119.54 | 132 | 1.10 | 125 | 1.05 | 71 | 0.59 | 103 | <u>0.87</u> | 72 | <u>0.60</u> | 46 | <u>0.39</u> | 40 | <u>0.33</u> | 26 | 0.22 | 13 | <u>0.11</u> | **628** | **5.25** |
| 8 | 122.49 | 124 | 1.01 | 118 | 0.96 | 107 | <u>0.87</u> | 90 | <u>0.74</u> | 44 | 0.36 | 45 | <u>0.37</u> | 42 | <u>0.34</u> | 30 | <u>0.25</u> | 11 | 0.09 | **611** | **4.99** |
| 9 | 95.79 | 89 | 0.93 | 101 | 1.05 | 93 | <u>0.97</u> | 64 | <u>0.68</u> | 70 | <u>0.73</u> | 27 | 0.29 | 13 | 0.14 | 20 | 0.21 | 6 | 0.06 | **483** | **5.04** |
| 10 | 84.56 | 98 | 1.16 | 55 | 0.65 | 66 | 0.78 | 39 | 0.47 | 27 | 0.32 | 16 | 0.19 | 58 | <u>0.69</u> | 12 | 0.14 | 5 | 0.06 | **376** | **4.45** |
| 11 | 118.47 | 130 | 1.10 | 113 | 0.95 | 64 | 0.54 | 48 | 0.41 | 49 | 0.41 | 27 | 0.23 | 37 | <u>0.31</u> | 31 | <u>0.26</u> | 14 | <u>0.12</u> | **513** | **4.33** |
| 12 | 104.02 | 71 | 0.68 | 77 | 0.74 | 55 | 0.53 | 40 | 0.39 | 53 | <u>0.51</u> | 22 | 0.21 | 18 | 0.17 | 33 | <u>0.32</u> | 13 | <u>0.12</u> | **382** | **3.67** |
| Total | **1536.9** | **1956** | **1.27** | **1767** | **1.16** | **1267** | **0.82** | **901** | **0.59** | **767** | **0.50** | **555** | **0.36** | **470** | **0.31** | **378** | **0.25** | **155** | **0.10** | **8216** | **5.35** |

a: the number of QTLs in each chromosome. b: average number of QTLs per cM. The average QTL numbers of the chromosomes larger than (or equal to) the total average QTL number are underlined.

